# SyRI: finding genomic rearrangements and local sequence differences from whole-genome assemblies

**DOI:** 10.1101/546622

**Authors:** Manish Goel, Hequan Sun, Wen-Biao Jiao, Korbinian Schneeberger

**Affiliations:** Max Planck Institute for Plant Breeding Research, D-50829 Cologne, Germany

**Keywords:** Genome comparison, structural rearrangements, structural variations, variant calling, genome alignments, genetics, genome assembly

## Abstract

Genomic differences range from single nucleotide differences to complex structural variation. Current methods typically annotate only differences in sequence (ranging from SNPs to large indels) but do not consider the full complexity of structural rearrangements (including inversions, translocations and duplications), where highly similar sequence changes in location, orientation, or copy-number. Here we present SyRI, a pairwise whole-genome comparison tool for chromosome-level assemblies. SyRI starts by finding rearranged regions and then searches for differences in the sequences, which are distinguished for residing in syntenic or rearranged regions. This distinction is important, as rearranged regions are inherited differently compared to syntenic regions.

## Background

Genomic differences form the basis for heritable phenotypic variations and their identification can allow us to decipher evolutionary past and gene function. Differences in genomes can range from single nucleotides to highly complex genomic rearrangements and are usually described as local sequence differences in comparison to a reference sequence. But even though the annotation of all sequence differences would be sufficient to reconstruct the actual sequence of a genome, local annotation of differences does not describe the full complex dependencies between the rearrangements. For example, a translocation is a genomic rearrangement where a specific sequence has moved from one region in the genome to another region. Although such a translocation could be described as a deletion at one region in the genome and an insertion at the other regions, this annotation would miss out on the fact that the deleted/inserted sequence is the same and that the sequence is not actually deleted but moved to a different region of the other genome. Like translocations, inversions and duplications also introduce such “differences in the structure” of a genome by introducing differences in location, orientation and/or copy number of the same sequence in the genome. But even though this information is usually not considered when analysing whole-genome sequencing data, such “differences in the structure” are relevant as they can be the basis for diseases phenotypes [1], reproductive strategies [2–4], and survival strategies [5].

Many of the state-of-the-art methods used to predict genomic differences utilize short or long read alignments against reference sequences [6]. Even though such alignments allow to find local sequence differences (like SNPs, indels and structural variation) with high accuracy, accurate prediction of differences in the structure remains challenging using read alignments alone. In contrast, whole-genome assemblies enable the identification of complex rearrangements as the assembled contigs are typically much longer and of higher quality as compared to raw sequence reads [7]. However, despite recent technological improvements to simplify the generation of whole-genome *de-novo* assemblies [8], there are so-far only a few tools which use whole-genome assemblies as the basis for the identification of genomic differences [9]. Available tools include AsmVar, which compares individual contigs of an assembly against a reference sequence and analyses alignment breakpoints to identify inversions and translocations [10], Assemblytics, which utilizes uniquely aligned regions within contig alignments to a reference sequence to identify various types of genomic differences including large indels or differences in local repeats [11] and Smartie-sv, which compares individual alignments between assembly and reference sequences [12].

Here, we introduce SyRI (Synteny and Rearrangement Identifier), a method to identify differences in structures as well as differences in the sequences (including small and complex local changes) of two whole-genome assemblies. SyRI expects whole-genome alignments (WGA) as input and starts by searching for differences in the structures of the genomes. Afterwards, SyRI identifies local sequence differences within both the rearranged and the non-rearranged (syntenic) regions. The distinction of local sequence differences in syntenic regions versus local sequence differences in rearranged regions is important as rearranged regions (and the variation in them) can show non-mendelian segregation patterns in offspring genomes. Finally, we validate SyRI’s performance with simulations and in comparison with existing tools used for the identification of genomic differences and apply SyRI to divergent genomes of five model species, including two *Arabidopsis thaliana* strains, for which we experimentally validate over 100 predicted translocations.

## Results

### The hierarchy in genomic differences

Genomes can differ in structure and sequence. *Differences in structure* occur if highly similar regions have different locations/orientations in different genomes (here we will refer to these regions as rearranged or non-syntenic regions). *Differences in sequence* are (simple to complex) differences in the nucleotide sequence within otherwise conserved regions. Importantly, differences in sequence can occur in both, rearranged as well as non-rearranged regions (Figure 1a). This introduces a hierarchy into the variation in genomes where some sequence differences reside in the conserved (syntenic) regions whereas other reside in rearranged regions of the genomes (e.g. a SNP within a translocated region).

**Figure 1:**
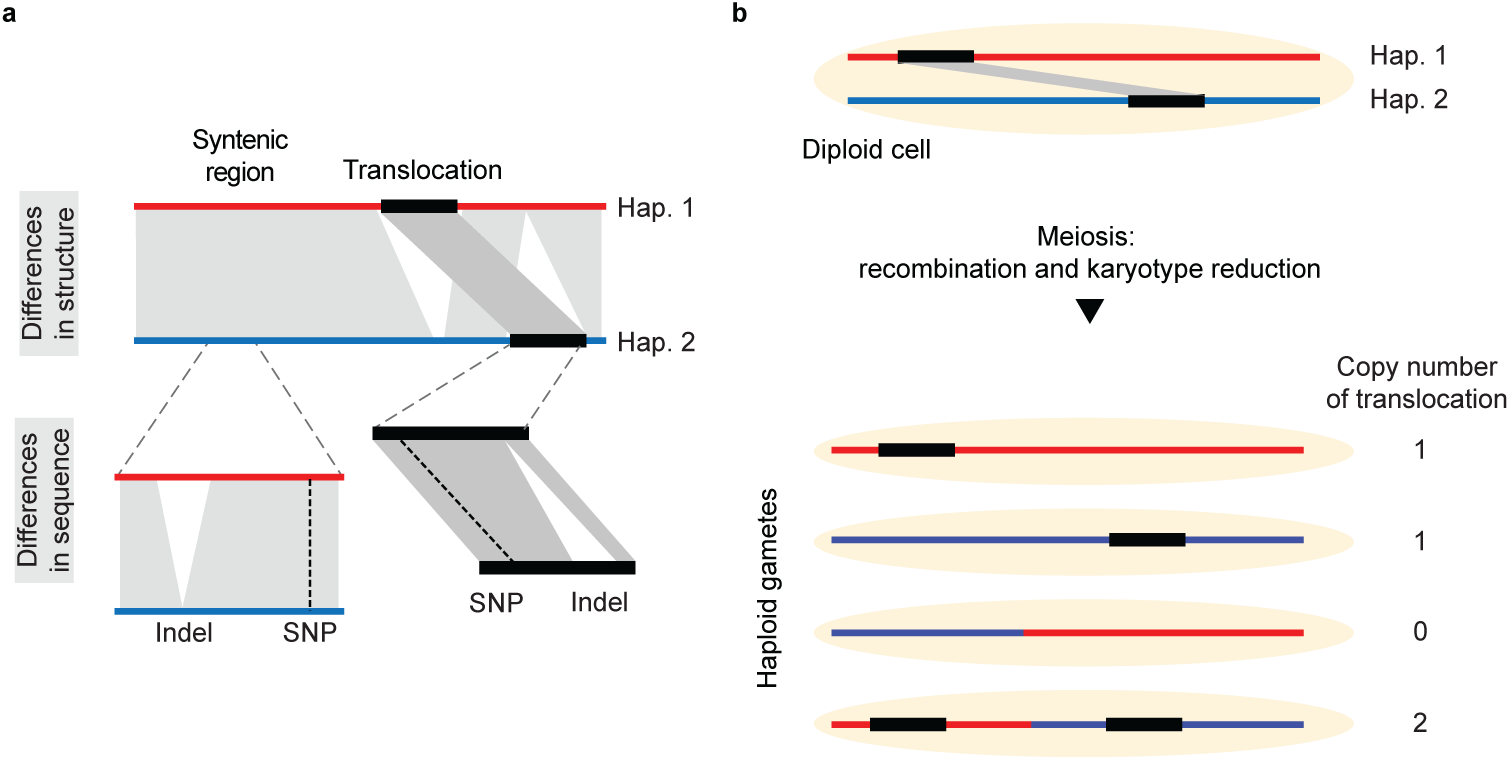
Hierarchy of genomic differences and their propagation. a) Genomic differences include differences in the structure (like inversions, translocations or duplications) as well as local sequence differences like SNPs/indels. Differences in sequence can occur in syntenic regions as well as in rearranged regions. b) A diploid cell containing two haplotypes, which can be distinguished by a translocation. Following meiosis and recombination, the haploid gametes can feature different copy-number variation for the translocated regions (and the sequence differences in it).

Even though reference-based resequencing analyses typically do not allow to distinguish between variation in syntenic versus non-syntenic regions, this distinction is important as non-syntenic regions (and the local variation in them) do not follow Mendelian segregation patterns in the offspring. Instead, due to the different locations in a genome, the inheritance of rearrangements can lead to changes in copy number or loss of the rearranged regions (Figure 1b).

### How SyRI works

SyRI is a whole-genome comparison tool that annotates all differences (including differences in structure and sequence) between two whole-genome assemblies (Figure 2). It starts by identifying all syntenic regions between the two genomes (Figure 2: Step 1). This at the same time also reveals all rearranged regions, as all regions which are not annotated as syntenic are rearranged by definition. In a second step, SyRI analyzes the location and orientation of the rearranged regions and groups them into inversions, translocations, and duplications (Figure 2: Step 2). As the last step, SyRI identifies sequence differences (like SNPs, indels, tandem duplications, copy-number changes etc) within both rearranged and syntenic regions (Figure 2: Step 3).

**Figure 2:**
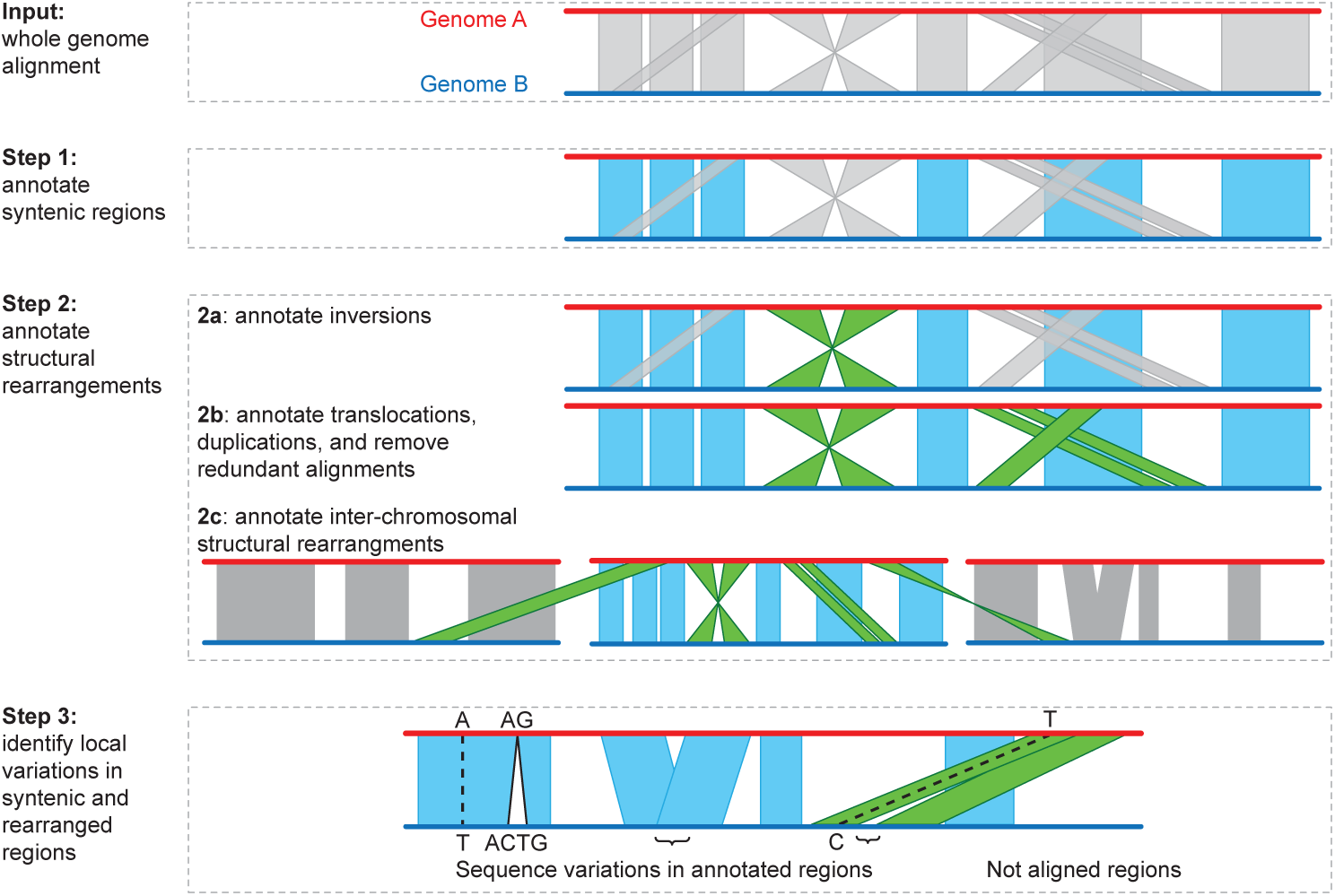
Workflow for the identification of genomic differences. SyRI uses whole-genome alignments (WGA) as input. A WGA consists of a set of local alignments, where each local alignment (grey polygon) connects a specific region in one genome to a specific region in the other genome. Step 1: SyRI identifies the highest scoring syntenic path between the corresponding genomes (blue alignments). The syntenic path represents the longest set of non-rearranged regions between two genomes. Step 2 (a-c): The remaining alignments are separated into structural rearrangements and redundant alignments. Structural rearrangements (green alignments) are classified into inversions, intra-chromosomal translocations and duplications, and finally inter-chromosomal rearrangements. Step 3: Local differences in the sequences are identified in all syntenic and rearranged regions. SNPs and small indels are parsed directly from the local alignments, whereas more complex sequence variation (e.g. like large indels and CNVs) are identified in the overlaps and gaps between consecutive local alignments. Also, all non-aligned regions in between syntenic and rearranged regions are reported for completeness.

To perform these three steps, SyRI generates different *genome graphs* which are generated from the local alignments from a pairwise whole-genome alignment (WGA). Here we used the MUMmer3 toolbox to perform WGA [13,14], but other alignments tools like minimap2 [15] can be used as well (Additional File 1: Note 1). In the following, we describe the individual steps of SyRI in more detail.

### Syntenic region identification

SyRI identifies syntenic regions by selecting the longest, non-contradicting subset of aligned regions which are syntenic to each other in the two genomes (Fig. 2: Step 1). For this, it selects all forward alignments between a pair of homologous chromosomes and generates a genome graph in the form of a directed acyclic graph (DAG) (Additional File 1: Note 2, Figure S1). SyRI then uses dynamic programming to identify the highest scoring path from the nodes that represent one end of a chromosome to the nodes that represent the other end (using similar algorithms as implemented in MUMmer [16,17]). This process is repeated for each pair of homologous chromosomes and after all syntenic regions are found, SyRI starts classifying the remaining non-syntenic regions into inversions, translocations, and duplications (Figure 2: Step 2).

### Inversion identification

An inversion is defined as a set of inverted alignments in-between two syntenic alignments (Additional File 1: Figure S2). Hence, if one genome would be reverse complemented then the alignments of an inversion would appear similar to syntenic alignments. Following this idea, SyRI selects all inverted alignments between a pair of corresponding chromosomes and reverse complements one of the chromosomes (Additional File 1: Figure S3, Note 3). Then, analogous to the syntenic path identification, SyRI again builds up a genome graph using these new forward alignments. From this graph, SyRI infers all possible candidate inversions between the two genomes (Additional File 1: Figure S3a). However, as candidate inversions can overlap and result in conflicting annotations (Additional File 1: Figure S3b), SyRI compares the annotations of all candidate inversions simultaneously and selects the best set of non-overlapping non-conflicting inversions while maximising the overall alignment score of the two genomes.

### Translocation and duplication (TD) identification

After synteny and inversion identification, all remaining alignments are either footprints of TDs or are redundant (repetitive) alignments. SyRI analyzes these alignments to find TDs while removing redundant alignments (Additional File 1: Note 4, Figure S4-S6). For this, SyRI first groups the alignments such that each group represents all alignments of a putatively rearranged region (candidate TD). Each candidate TD is given a score based on its alignment length and gap length between consecutive alignments. Low scoring candidates and those that are overlapping with syntenic or inverted regions are filtered out.

As a result of repeats, rearranged regions can have different candidate TDs aligning to different copies of the same repeat region. Therefore, overlapping candidate TDs often result in conflicting annotations. SyRI resolves these overlapping candidate TDs by selecting the non-conflicting subset of candidate TDs with the highest alignment score (Additional File 1: Note 4, Figure S5, S7).

### Grouping of alignments to generate annotation blocks

As a final step of the analysis of differences in structure, SyRI combines all neighbouring alignments of the same type to form annotation blocks. For example, a syntenic block would contain all consecutive uninterrupted syntenic alignments. Likewise, inversion or TD blocks include all alignments which together form the extent of an inversion or of a TD.

### Identification of sequence differences

Finally, SyRI reports differences in the sequence. This includes small variation (like SNPs and small indels) which are found in the local alignments generated by the whole-genome alignment algorithm as well as larger structural variations (like indels or CNVs), which are not part of the local alignments (Figure 2: Step 3). To find these structural variations, SyRI analyses the gaps between all consecutive alignments in annotation blocks and identifies indels, highly divergent regions (HDRs) and CNVs/tandem repeats (Additional File 1: Figure S8) similar to the SV identification of Assemblytics [11]. Finally, SyRI reports all *un-aligned regions* which are not part of any annotation block.

### Performance evaluation using simulated genomes

We validated SyRI using a simulation study with 400 simulated genomes which were generated by randomly inserting inversions, translocations, duplications, and indels into the reference sequence of *A. thaliana* (Methods). Comparing these genomes against the unaltered reference sequence, SyRI showed sensitivity and precision values of more than 95% for each of the different types of rearrangements (Figure 3a, b). False results were usually a consequence of rearranged regions in repeat regions where alternative annotations were equally likely.

**Figure 3:**
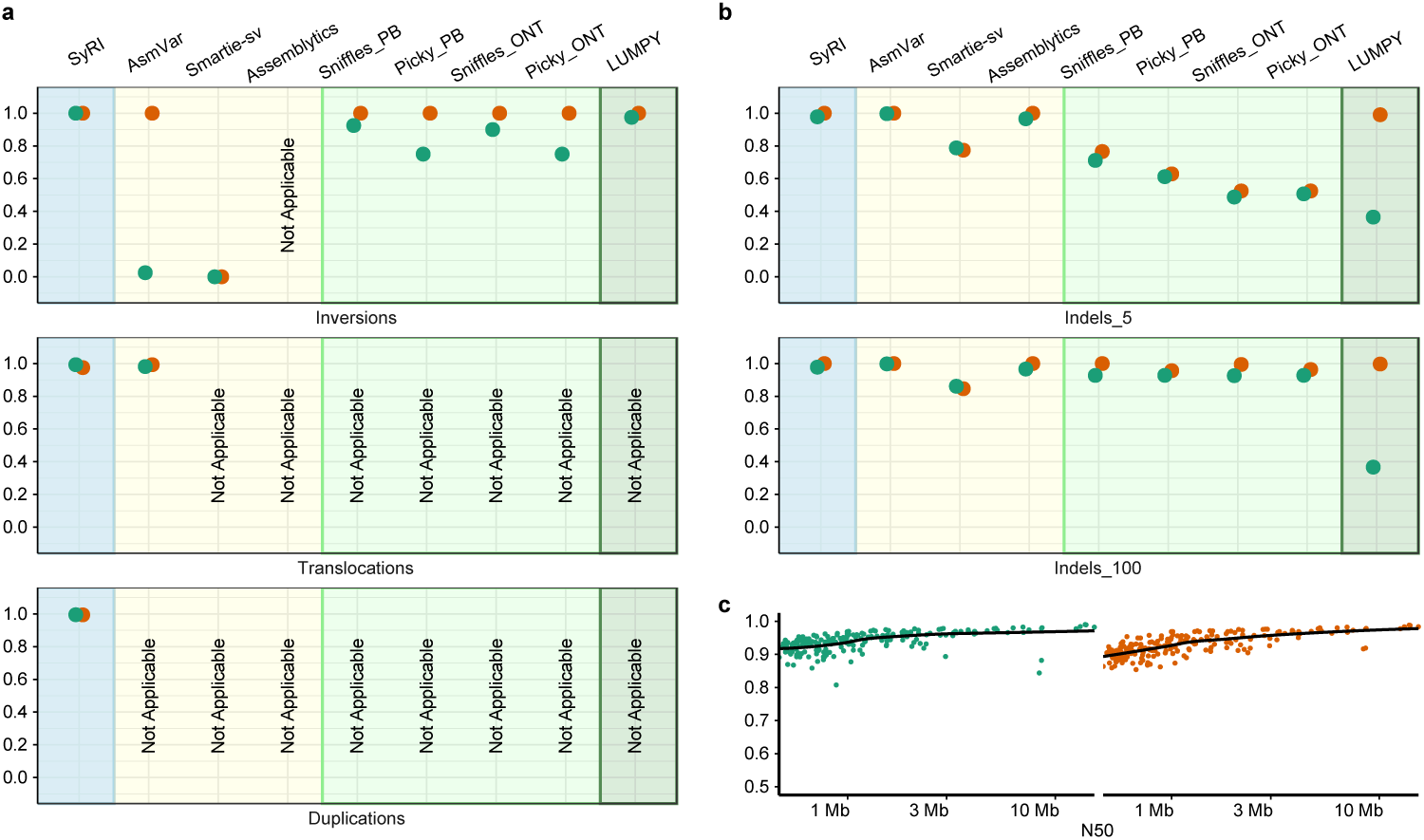
SyRI’s performance compared to six other tools using simulated assemblies and on incomplete assemblies. Sensitivity (green points) and precision (orange points) values are shown for the identification of (a) rearranged regions and (b) indels. Each point describes the median sensitivity (and precision) values from the analysis of 100 simulated genomes. Background colors represent the data type required by the respective tools: light blue for chromosome-level *de novo* assembly, pale yellow for *de novo* assembly, pale green for long sequencing reads (both PacBio (PB) and Oxford Nanopore (ONT) reads), green for short sequencing reads. (a) Performance values for inversions, translocations and duplications. (As AsmVar found inversions in only 40 samples, performance values for AsmVar considered only these 40 samples.) (b) Performance values for indel identification using stringent (upper) or relaxed (lower) thresholds for matching predicted vs. simulated indel positions. (LUMPY cannot identify insertions, however, sensitivity calculation did not account for that.) (c) SyRI’s performance for the identification of rearranged regions from incomplete assemblies. Each point represents the performance value generated with one of the simulated assemblies. The black line represents the polynomial fit.

We compared this to the performance of six other tools: AsmVar, smartie-sv, assemblytics, sniffles, picky, LUMPY [10–12,18–20]. AsmVar, smartie-sv, and assemblytics identify genomic differences using whole-genome assemblies, sniffles and picky are based on long read alignments, and LUMPY uses short read alignments against a reference sequence. For the tools that require sequencing read data, we simulated reads from the whole genome assemblies and aligned them to the reference sequence (Methods). However, unlike SyRI, these tools are not designed to find all genomic differences (Additional File 2: Table S1). Generally, for variant callers “translocation” identification is limited to identification of chromosome-end rearrangements whereas “duplication” identification is limited to identification of tandem duplications, in contrast, SyRI identifies all genome-wide rearrangements.

In our simulations, SyRI outperformed all other tools by consistently identifying all genomic differences (Figure 3a, b). All tools identified indels with high sensitivity and precision, however, only SyRI, AsmVar and Assemblytics identified indel breakpoints with high accuracy. For inversions, only LUMPY and sniffles performed comparable to SyRI, whereas, for translocations, SyRI and AsmVar showed the best performances.

### Performance evaluation using real genomes

To test SyRI’s performance with real data, we applied it to identify the variation in the human genome NA19240 for which gold standard variation data was recently published (Additional File 1: Figure S9, Methods) [21]. These gold standard variation data include differences which were predicted based on whole-genome shotgun read alignments against the reference sequence and therefore, include variation from both haplotypes of this genome. In addition, a whole-genome assembly, which includes only one of the two haplotypes, is available as well [22].

Using this whole genome assembly in comparison to the reference sequence, SyRI identified 55.2% (9685 out of 17545) of the gold standard insertions, 54.5% (9494 out of 17391) of the deletions, and 49.7% (81 out of 163) of the inversions (Additional File 1: Figure S10, Methods), which is consistent with the presence of only one of the haplotypes in the assembly. In comparison to the other tools tested here, SyRI identified a higher proportion of all different types of genomic variation of the gold standard variation (Additional File 1: Figure S10).

For a second comparison, we generated a chromosome-level assembly of the (homozygous) *A. thaliana* Ler genome using long PacBio reads. The assembly CN50 and CL50 values (chromosome number normalized N50 and L50 values) were 12.6 Mb and 1 respectively (Additional File 2: Table S2, Methods, Additional File 1: Figure S11) [23].

We again applied the other tools to identify differences between the Col-0 and Ler genomes (Additional File 1: Figure S12, Methods). SyRI and AsmVar found similar numbers of indels, while Assemblytics found a much higher number of indels. However, all of these assembly-based methods identified more indels than the read based methods. More inversions were identified in the read based methods compared to SyRI and AsmVar. A similar number of translocations were identified by SyRI and AsmVar, whereas only SyRI identified duplications. The performance of the read based methods was limited by severe overcalling of genomic differences. For every read based method, the cumulative length of deletions was more than 150Mbp, with multiple large deletions (>1,000,000, Additional File 2: Table S3) while the genome size of *A. thaliana* is ∼135Mbp [24]. Moreover, two of the tools annotated more than 70Mbp and 14Mbp of inversions, respectively. These results are highly improbable and most likely resulted from the incorrect classification of structural rearrangements.

### Effect of genome contiguity

SyRI requires whole-genome alignments from chromosome-level assemblies as input. If one or both of the assemblies is/are incomplete (i.e. not on chromosome level), pseudo-chromosomes can be generated using homology between the assemblies themselves or using homology to a chromosome-level reference sequence, which can be performed with tools like RaGOO (Additional File 1: Note 5, preprint [25]). To analyze the effect of the contiguity of the original assembly on SyRI’s performance, we performed a simulation analysis where we first generated multiple incomplete assemblies from the chromosome-level assembly of *A. thaliana* Ler by randomly breaking the chromosome-level scaffolds in unconnected pieces (Methods). These scattered assemblies were then reassembled with RaGOO using their homology to the *A. thaliana* Col-0 reference genome (preprint [25]).

We then identified rearranged regions in each of these re-assemblies by comparing them to the reference sequence using SyRI. These resulting rearrangements were then compared with the results SyRI generated when comparing the original chromosome-level assembly of Ler against the reference sequence.

More than 90% of the assemblies with N50 of more than 470 Kb (before the homology based reassembly) had a sensitivity of more than 0.9 (Figure 3c). Similarly, more than 90% of the assemblies with N50 more than 674Kb had a precision of more than 0.9. The shortest assemblies we generated had N50 values in the range of 470-500 Kb and the predictions based on these assemblies still had average sensitivity and precision values of 0.92 and 0.90 respectively.

We then evaluated SyRI’s efficiency in identifying rearranged regions when both genomes are at scaffold level. For this, we generated scattered assemblies from both the Col-0 reference sequence and the Ler assembly. Since current pseudo-chromosome generation tools only concatenate scaffolds of one assembly using homology with another assembly, we developed a heuristic script to generate homology-based pseudo-chromosomes using two incomplete assemblies (Additional File 1: Note 5). As before, we identified rearranged regions from these pseudo-genomes and compared them to the rearranged regions identified between the full-length assemblies. For assemblies with N50 of more than 868Kb and 721Kb, sensitivity and precision values of more than 0.7 in more than 70% of the cases (Additional File 1: Figure S13). For assemblies with lower contiguity (N50: 470-500 Kb), sensitivity and precision were 0.56 and 0.65, respectively.

Together this shows that the prediction of genomic rearrangements is nearly complete even if one of the genomes is not on chromosome-level, but has assembly contiguity of N50 > 500 kb. If both assemblies are not on chromosome-level the quality of the predictions is reduced, however, it is still possible to get useful insights on a subset of the rearrangements.

### Runtime estimation when comparing human, yeast, fruit fly, and maize genomes

To analyze SyRI’s runtime performance, we searched for intra-species genomic differences in four different model organisms: human, yeast, fruit fly, and maize (Additional File 2: Table S2). For its application to human genomes, we compared whole-genome assemblies of NA12878 and NA19240 against the reference genome GRCh38.p12 [22,26,27]. For yeast, we compared the *de novo* assembly of strain YJM1447 against the reference genome from strain S288C [28,29]. For fruit fly (*Drosophila melanogaster*), the *de novo* assembly of strain A4 was compared to the reference genome [30,31]. For maize, we compared the *de novo* assembly of PH207 against the B73 reference genome [32,33]. To limit computational requirements, we masked the highly repetitive maize genome while all other genomes were analysed without masking [34].

SyRI was used to predict syntenic and rearranged regions as well as local sequence differences in all genomes (Table 1, Additional File 1: Figure S14–S18). In each comparison, including human, at least 5% of the assembled genomes were found to be non-syntenic. The CPU runtime for the smaller and simpler yeast genomes was 34.5 seconds, whereas for the two human genomes SyRI took ∼10 minutes, while memory usage was less than 1 GB for each of the comparisons (Table 1) (without considering SNPs and small indels parsing). The exception was the comparison of the repetitive maize genomes, which took ∼1hr of CPU time and ∼6GB of RAM. Since SyRI considers all alignment combinations, the runtime and memory usage can be high in repetitive genomes (Additional File 1: Note 6 and Figure S19). However, the number of alignments can be drastically reduced by decreasing the WGA sensitivity (i.e. omitting small, 10-100s bp alignments), which in turn decreases runtime and memory consumption of SyRI.

**Table 1:**
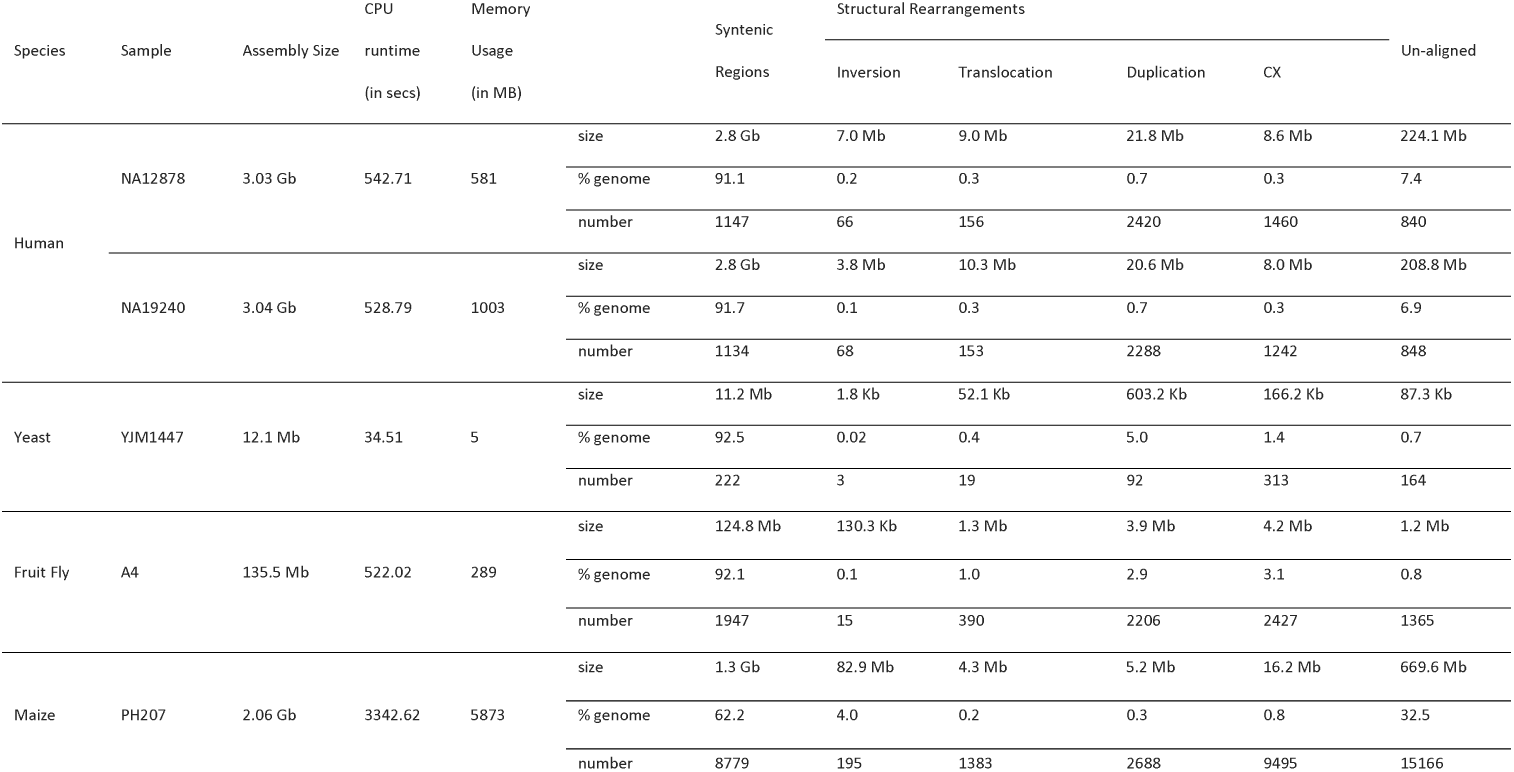
Structural differences identified by SyRI and corresponding computation resources.

### Experimental validation

To validate some of the predicted translocations in the genome of *A. thaliana* L*er*, we used a genetic approach which was based on the observation that recombinant offspring genomes feature different copy numbers of translocated DNA (Figure 1b; 4a), while non-translocated regions always occur with the same copy number. The actual copy number of translocated DNA in a recombinant genome relies on the genotypes at the two insertion sites of the translocation. For example, translocated DNA is duplicated if the two insertion sites of a translocation are combined into one recombinant haplotype.

We used available whole-genome sequencing data of a set of 50 F_2_ recombinant plants, which were generated by crossing Col-0 and L*er*, followed by self-pollination of the resulting F1 hybrids (preprint [35]). We aligned the short reads (∼5x genome coverage/sample) to the Col-0 reference sequence and used the genotypes at ∼500k SNP markers to reconstruct the parental haplotypes using TIGER (Figure 4b) [36,37].

**Figure 4:**
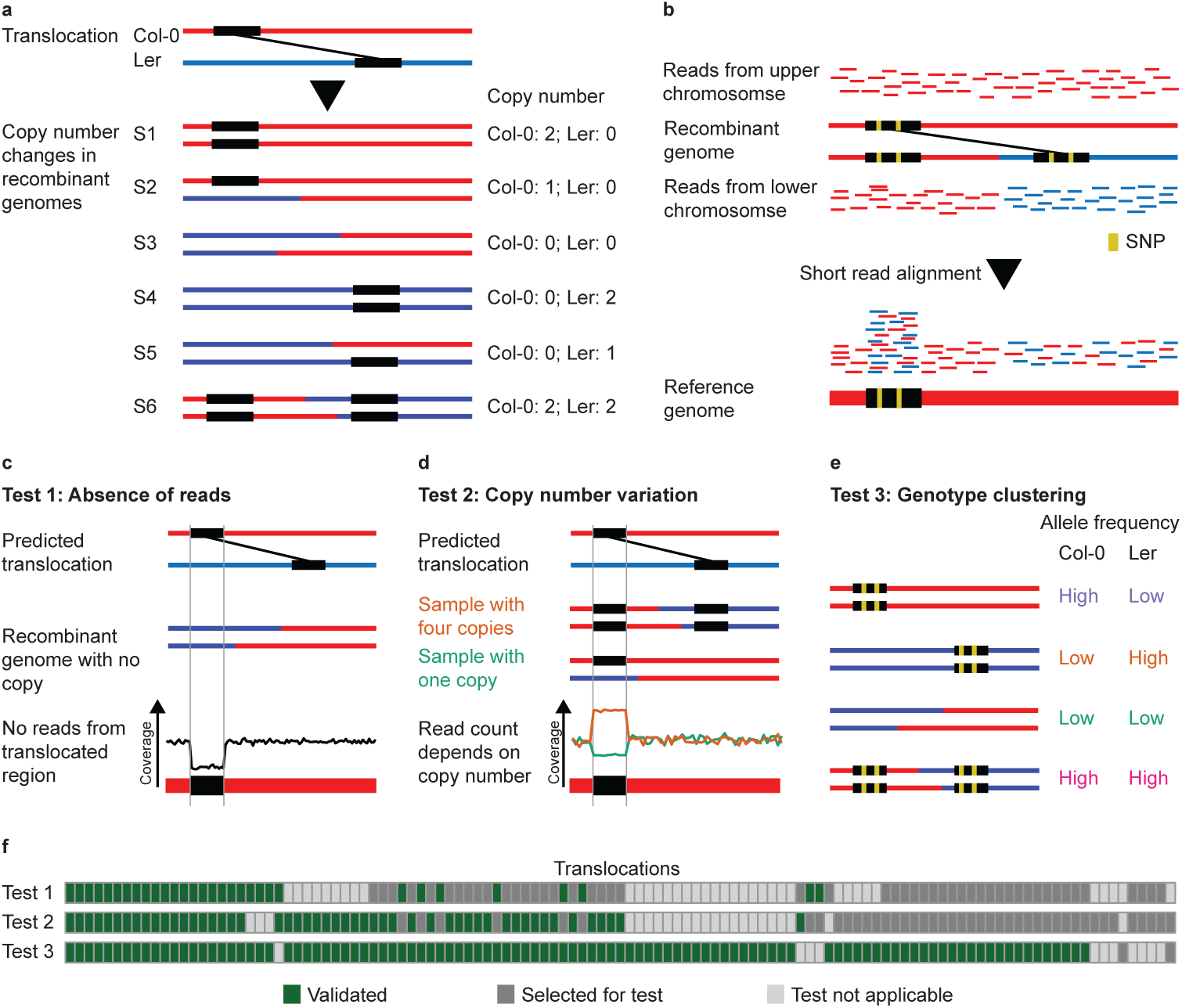
Recombination introduces copy-number variation. (a) Recombination between two haplotypes with translocated regions can lead to copy-number differences in the recombined genomes. (b) Such differences can be observed by aligning short-read sequencing data from recombinant genomes to the reference genome. (c-e) Three different tests to assess the existence of the predicted translocations have been applied. These included (c) testing for the absence of reads in samples with no copy of the translocated DNA, (d) goodness-of-fit between expected copy number and observed copy-number, and (e) clustering of samples with the same genotypes at the translocation. (f) In the heatmap, columns correspond to individual translocations and rows correspond to the three different tests, while the colour of a cell represents whether a translocation was validated (green), was selected but could not be validated (dark grey), or was filtered out as the test was not applicable (grey).

Based on this haplotype information, we estimated the expected copy number for 117 translocations, which were larger than 1kb, in each of the samples. The expected copy-number was then compared to the outcome of three different tests. The first two tests were based on the assumption that all reads from a translocated region align to the same loci in the reference genome independent of the actual location of the rearranged region in the sequenced sample (Figure 4b) [38]. This allows estimating copy-number of a translocation using read coverage in the respective region of the reference. For the first test, we analysed the absence of reads in translocated regions in recombinant genomes, which were predicted to feature no copy of the translocated region (Figure 4c) (using 0.2x read coverage as a cut-off to distinguish between absence or presence of a translocation). For the second test, we assessed the goodness-of-fit between expected copy-number and observed copy-number for a translocation across all recombinants (as estimated from the normalized read counts in the translocation regions; Figure 4d; Methods). The third test was based on the sequence differences between the different alleles of a translocation. For this, we tested differences in the read counts supporting either the Col-0 (or L*er*) alleles of a translocation. Depending on the copy number of the different alleles of a translocation, the allele count should also vary. In consequence, samples with the same genotypes at the two loci of a translocation should have similar allele counts, whereas samples with different genotypes should also show different allele counts (Figure 4e; Methods).

Out of 117 translocations, 108 (92.3%) could be confirmed by at least one test (Figure 4f). We manually checked the read alignments of the nine translocations that could not be confirmed and found support for the existence of each of the translocations, which however had not been strong enough to be identified by any of the three test criteria. In summary, this supports that a large majority of the translocations predicted by SyRI are real.

## Discussion

We introduced SyRI, a tool that identifies genomic differences between two whole-genome assemblies. The genomic differences include structural differences as well as differences in sequences. However, instead of identifying differences directly, SyRI starts by identifying all syntenic regions between the genomes, as all other (non-syntenic) regions are rearranged by definition.

Once the structural differences are found, SyRI identifies local sequence differences in both syntenic and rearranged regions. The identification of local sequence differences in rearranged regions introduces a hierarchy of genomic variations (e.g., SNPs in translocated regions). This distinction is important as rearranged regions are differently inherited as compared to syntenic regions. If this is not accounted for genotypes in rearranged SNPs can confound the interpretation of genomic patterns during selection screens, genome-wide association or recombination analysis [39,40]. SyRI now offers a straight-forward solution to filter SNPs in rearranged regions assuming whole-genome assemblies are available.

SyRI provides comprehensive and accurate genomic difference annotation using chromosome-level assemblies. Even though generating *de novo* assembly is still not trivial, consistent improvements are being made. Finally, though SyRI is based on a genome graph that is build up from the local alignments of a WGA, this algorithm can be easily adapted for rearrangement identification in other types of genome graphs as well [41,42].

## Conclusions

We have developed SyRI which, to our knowledge, is the first tool to identify all structural and sequence differences between two genome assemblies. This novel approach is highly efficient and includes a classification of sequence differences for being in syntenic or rearranged regions. Using SyRI, we identified genomic rearrangements and sequence differences in humans, *A. thaliana*, fruit fly, yeast, and maize genomes. Additionally, we validated the existence of more than 100 predicted translocations. SyRI is available as an open source tool and is being actively developed and improved.

## Methods

### Long read sequencing of the genome of *A. thaliana* L*er*

*A. thaliana* L*er* plants were grown in the greenhouse at the Max Planck Institute for Plant Breeding Research. DNA was extracted using the NucleoSpin^®^ Plant II Maxi Kit from Macherey-Nagel. We used the PacBio template prep kit > 20 kb for Sequel systems (SMRTbell Template Prep Kit 1.0-SPv3) with damage repair (SMRTbell Damage Repair Kit -SPv3) and BluePippin size selection for fragments > 9/10 kb. Sequencing of two SMRT cells was done with the Sequel Sequencing Plate 1.2 and the Sequel Binding Kit 1.0. Movie Time 360 min.

### Assembly generation

Raw reads were filtered to remove small and low-quality reads (length<50bp and QV<80), corrected and *de novo* assembled using Falcon [43], followed by polishing with Arrow in the SMRTLink5 package, and finally corrected using Illumina short read alignments with reads from an earlier project [44]. The contigs from organellar DNA sequences were removed, all others were anchored into pseudo-chromosome based on homology with the reference sequence. Adjacent contigs were connected with a stretch of 500 “N” characters.

### Whole-genome alignments

All assemblies used in this work were filtered to select only chromosome-representing contigs (unplaced scaffolds were removed). We used the *nucmer* alignment tool from the MUMmer toolbox [14] to perform WGAs. Nucmer was run with --maxmatch to get all alignments between two genomes including -c, -b, and -l parameters which were selected to balance alignment resolution and runtime based on genome size and number of repeat regions (full commands are available in Additional File 2: Table S4). Alignments were filtered using the *delta-filter* tool and the filtered delta files were converted to the tab-delimited files using the *show-coords* command. Before whole-genome alignments, both maize genomes were masked using RepeatMasker v4.0.6 [45].

### Simulating rearranged genomes

We simulated structural rearrangements in the *A. thaliana* reference genome using the *R* package *RSVSim* [46]. We simulated 40, 436, and 1241 events for inversions, translocations, and duplications respectively and for each rearrangement 100 genomes were simulated. The number of rearrangements and their corresponding sizes were sampled from real differences found between the Col-0 and Ler genomes. For simulating indels, we used SURVIVOR [47] to simulate 100 genomes containing 1,000 indels in the range of 1-500 bps.

From these rearranged genomes, we simulated PacBio and Nanopore reads using SURVIVOR. We used the *A. thaliana* long read data generated by Michael et al. (NCBI project accession: PRJEB21270) to generate read profiles required by SURVIVOR and simulated reads to get a 30x coverage [48]. Short reads were simulated using wgsim (parameters used: -e 0.001 -d 550 -N 12000000 -1 150 -2 150) to get 30x coverage [49]. All reads were aligned to the *A. thaliana* reference genome using minimap2 and the alignments were converted from SAM to BAM format and sorted using samtools [15,50].

### Running tools on simulated genomes

SyRI: Genome assemblies were aligned using nucmer (Additional File 2: Table S4) and SyRI was run with default parameters. Assemblytics: We used the same alignments generated by nucmer as used for SyRI. The default value for unique sequence length was used and variants size was set from 1 to 100,000bp. AsmVar: The tool was run based on the demo script provided with the tool. For genome alignment, lastdb was run using the default parameters, whereas lastal and last-split were run using the parameters provided in the above demo [51]. Similarly, variants were detected using ASV_VariantDetector tool of AsmVar with the default parameters. Smartie-sv: The pipeline was run using the default settings. However, the number of jobs to be run in parallel and job wait time was adjusted to make it suitable for the computer resources available. Sniffles: Sniffles was run separately for PacBio and Nanopore simulated reads using the default parameters. Alignments were generated through minimap2 and converted to BAM and sorted using samtools. Picky: Picky was run using the same methodology and parameters as described by the authors for both PacBio and Nanopore reads. LUMPY: Reads were aligned by minimap2, and the alignments were pre-processed using samblaster [52] and samtools as per the instructions provided by the authors. While running LUMPY, paired-end read distribution parameters were changed to match the simulated reads (mean:550, read_length:150, min_non_overlap:150).

Structural rearrangements predicted by these tools were considered to match an original simulated rearrangement if the end-points for the predicted rearrangement were within ±150bp of the end-points of the original rearrangement. For indels, we compared the location and size of the predicted indels with the simulated indels, allowing for error in both location and size. Two different error limits were checked: 5 and 100 bp.

### Performance evaluation with real genome data

For both the *A. thaliana* (L*er*) and the human (NA19240) genome, we used the same methods as above to simulate sequencing reads from whole genome assemblies, to perform alignments with the reference genomes, and to identify genomic differences. For human genomes, we used the error profiles provided by SURVIVOR [47]. Number and sizes of the variations were extracted from the output files using in-house scripts. For the AsmVar comparison of Col-0 vs Ler, we used the .svd output file instead of the .vcf output file as the former had better suited annotations for our analysis. An indel was considered as identified if there was a simulated indel of the same type (insertion or deletion) within 100bp of the location of the predicted indel and the size difference between two indels was not more than 100bps.

### Comparison with the gold standard variation dataset

Variant calls for the gold standard dataset were downloaded from the NCBI [21]. The variants were generated with an older version human reference genome (GRCh38) and were therefore re-mapped to the newer GRCh38.p12 version of the human reference genome using the NCBI Genome Remapping Service. An indel from the gold standard dataset was considered to be identified if a predicted indel of the corresponding type existed within the surrounding 100bp. For inversion predictions, we checked the overlap between inversions from the gold dataset and the inversions, inverted translocations, and inverted duplications as annotated by SyRI.

### Pseudo-chromosome generation and output comparison

We generated 200 fragmented assemblies of the L*er* genome by introducing 10-400 random breakpoints. Pseudo-genomes were generated for each of the fragmented assemblies using RaGOO with default parameters. Additionally, we generated 100 fragmented assemblies each of Col-0 and L*er* again by introducing 10-400 random breakpoints. These fragmented assemblies were assembled by a heuristic script (Additional File 1: Note 5) to generate pseudo-molecules. For 16 assemblies, pseudo-molecule generation failed and these samples were skipped from further analysis. A genomic rearrangement identified from the pseudo-genomes was considered to be correct if the same rearrangement type was present within 100bp up or downstream.

### Data extraction and transformation of the 50 recombinant genomes

For validation, we used whole-genome sequencing data of 50 F_2_ recombinant plants that we generated recently (preprint [35]). We extracted allele count information from consensus call files generated by SHORE [53]. For each predicted translocation, we estimated its copy number as the ratio between average read-coverage for the translocated region and the average read-coverage across the entire genome of the respective sample. Translocations in the centromeric regions and for which more than 25% of the translocated sequence had at least 10% reads with Ns were filtered out. For allele count analysis, we selected high-confidence (25bp conserved in both directions) SNPs in translocated regions as markers.

### Validation of translocations: Absence of reads (Test 1)

We selected F2 samples which, according to predicted genotypes, should have lost the translocated DNA and thus should not give rise to any reads from the translocated region. Only translocations for which at least two samples that had lost the translocated regions existed were tested. And only those translocations for which all tested samples had no reads were considered as validated.

### Validation of translocations: Expected vs. observed copy number (Test 2)

For each translocation, we selected samples which had different genotypes at the two associated loci for the translocation. This removes some of the samples with two copies and helps to remove a bias towards genomes with a copy number of two, which can affect this test. We further selected translocations for which we found samples with at least three different copy-number values predicted. A linear model was fit using the *lm* function in R. P-values for the model-fit were adjusted for multiple testing using the *BH-*method [54], and translocations for which adjusted *p*-values were less than 10^-6^ and slope more than 0.75 were considered as valid.

### Validation of translocations: Genotype clustering (Test 3)

Allele count values at the SNP markers were normalized and outliers (markers having very high allele counts) were removed. Translocations were tested only when they had at least two different classes of samples (genotypes) with each class having at least three samples, and at least three SNP markers in the translocated regions. Translocations for which alternate allele counts did not change across the samples (variance < 1) were also filtered out.

#### Cluster fit calculation

First, the distance between two samples was defined as the Euclidean distance between their reference allele counts and alternate allele counts. Then, the *closeness_score* was calculated as the sum of ratios of the average distance between the samples belonging to a genotype to the average distance to samples of other genotypes.

#### Simulating distributions

Background distributions for the closeness_score were simulated by generating random clusters. For each sample, allele counts (reference and alternate) were sampled using a Poisson distribution. For true translocations, the closeness_score would be low as samples from the same genotype would be much closer to each other, whereas samples from different genotypes would be far. For each translocation, we calculated the lower-tail *p*-value of retrieving the corresponding *closeness_score. P*-values were adjusted for multiple testing using *BH*-method, and translocations with *p*-value < 0.05 were considered valid.

## Declarations

### Availability of data and materials

The assembly of the L*er* genome has been submitted to the European Nucleotide Archive (http://www.ebi.ac.uk) and is publicly available under the accession number GCA_900660825. All other assemblies are publicly available at NCBI (https://www.ncbi.nlm.nih.gov/), and their accession numbers are GCA_000001735.3 [55], GCA_000001405.27 [26], GCA_002077035.3 [27], GCA_001524155.4 [22], GCA_000146045.2 [29], GCA_000977955.2 [28], GCA_000001215.4 [31], GCA_002300595.1 [30], GCA_000005005.6 [33], GCA_002237485.1 [32]. Further details about the assemblies are in Additional File 2: Table S2. BAM files for the 50 F_2_ recombinant genomes are available at European Nucleotide Archive under the project ID PRJEB29265 (preprint [35]). SyRI is freely available under the MIT license and is available online [56]. SyRI is developed using Python3 and is platform independent.

## Authors’ contributions

The project was conceived by KS and WBJ. MG and KS developed the algorithms. MG implemented SyRI and performed all analyses. HS processed recombinant genome sequencing data and identified crossing-over sites. WBJ generated the L*er* assembly. The manuscript was written by MG and KS with inputs from HS and WBJ. All authors read and approved the final manuscript.

## Acknowledgements

The authors would like to thank Ulrike Hümann for help with plant work and Detlef Weigel and Gunnar Klau for helpful comments on the manuscript. We would also like to thank the researchers at the Genome Institute at Washington University School of Medicine who shared the assemblies for the NA12878 and NA19240 genomes.

## List of Additional Files

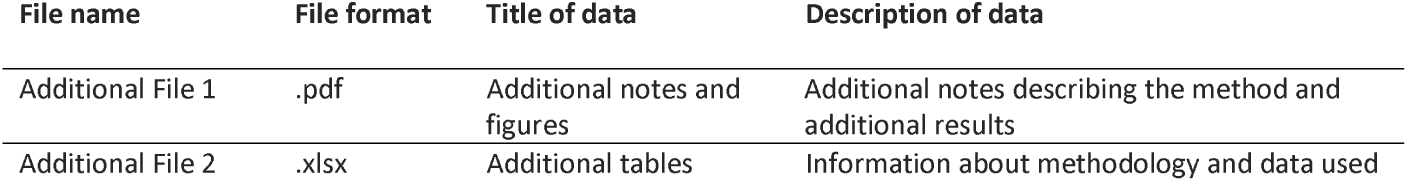

**List of Additional Files:**
| File name | File format | Title of data | Description of data |
| --- | --- | --- | --- |
| Additional File 1 | .pdf | Additional notes and figures | Additional notes describing the method and additional results |
| Additional File 2 | .xlsx | Additional tables | Information about methodology and data used |

## References

1. Weischenfeldt J, Symmons O, Spitz F, Korbel JO. Phenotypic impact of genomic structural variation: insights from and for human disease. Nat Rev Genet. 2013; 14(2):125–38.

2. Tuttle EM, Bergland AO, Korody ML, Brewer MS, Newhouse DJ, Minx P, et al. Divergence and Functional Degradation of a Sex Chromosome-like Supergene. Curr Biol. 2016; 26(3):344–50.

3. Küpper C, Stocks M, Risse JE, dos Remedios N, Farrell LL, McRae SB, et al. A supergene determines highly divergent male reproductive morphs in the ruff. Nat Genet. 2016; 48(1):79–83.

4. Lamichhaney S, Fan G, Widemo F, Gunnarsson U, Thalmann DS, Hoeppner MP, et al. Structural genomic changes underlie alternative reproductive strategies in the ruff (Philomachus pugnax). Nat Genet. 2016; 48(1):84–8.

5. Lowry DB, Willis JH. A Widespread Chromosomal Inversion Polymorphism Contributes to a Major Life-History Transition, Local Adaptation, and Reproductive Isolation. Barton NH, editor. PLoS Biol. 2010; 8(9):e1000500.

6. Guan P, Sung W-K. Structural variation detection using next-generation sequencing data. Methods. 2016; 102:36–49.

7. Simpson JT, Pop M. The Theory and Practice of Genome Sequence Assembly. Annu Rev Genomics Hum Genet. 2015; 16(1):153–72.

8. Jiao W-B. The impact of third generation genomic technologies on plant genome assembly. Curr Opin Plant Biol. 2017; 36:64–70.

9. Sedlazeck FJ, Lee H, Darby CA, Schatz MC. Piercing the dark matter: Bioinformatics of long-range sequencing and mapping. Nat Rev Genet. 2018; 19(6):329–46.

10. Liu S, Huang S, Rao J, Ye W, Krogh A, Wang J. Discovery, genotyping and characterization of structural variation and novel sequence at single nucleotide resolution from de novo genome assemblies on a population scale. Gigascience. 2015; 4(1).

11. Nattestad M, Schatz MC. Assemblytics: a web analytics tool for the detection of variants from an assembly. Bioinformatics. 2016; 32(19):3021–3.

12. Kronenberg ZN, Fiddes IT, Gordon D, Murali S, Cantsilieris S, Meyerson OS, et al. Highresolution comparative analysis of great ape genomes. Science (80-). 2018; 360(6393):eaar6343.

13. Marçais G, Delcher AL, Phillippy AM, Coston R, Salzberg SL, Zimin A. MUMmer4: A fast and versatile genome alignment system. Darling AE, editor. PLOS Comput Biol. 2018; 14(1):e1005944.

14. Kurtz S, Phillippy A, Delcher AL, Smoot M, Shumway M, Antonescu C, et al. Versatile and open software for comparing large genomes. Genome Biol. 2004; 5(2):R12.

15. Li H. Minimap2: pairwise alignment for nucleotide sequences. Birol I, editor. Bioinformatics. 2018; 34(18):3094–100.

16. Delcher AL, Phillippy A, Carlton J, Salzberg SL. Fast algorithms for large-scale genome alignment and comparison. Nucleic Acids Res. 2002; 30(11):2478–83.

17. Delcher AL, Kasif S, Fleischmann RD, Peterson J, White O, Salzberg SL. Alignment of whole genomes. Nucleic Acids Res. 1999; 27(11):2369–76.

18. Sedlazeck FJ, Rescheneder P, Smolka M, Fang H, Nattestad M, von Haeseler A, et al. Accurate detection of complex structural variations using single-molecule sequencing. Nat Methods. 2018; 15(6):461–8.

19. Gong L, Wong C-H, Cheng W-C, Tjong H, Menghi F, Ngan CY, et al. Picky comprehensively detects high-resolution structural variants in nanopore long reads. Nat Methods. 2018; 15(6):455–60.

20. Layer RM, Chiang C, Quinlan AR, Hall IM. LUMPY: a probabilistic framework for structural variant discovery. Genome Biol. 2014; 15(6):R84.

21. Chaisson MJP, Sanders AD, Zhao X, Malhotra A, Porubsky D, Rausch T, et al. Multiplatform discovery of haplotype-resolved structural variation in human genomes. Nat Commun. 2019; 10(1):1784.

22. The Genome Institute at Washington University School of Medicine. NA19240_prelim_3.0. https://www.ncbi.nlm.nih.gov/assembly/GCA_001524155.4. 2017.

23. Jiao W-B, Accinelli GG, Hartwig B, Kiefer C, Baker D, Severing E, et al. Improving and correcting the contiguity of long-read genome assemblies of three plant species using optical mapping and chromosome conformation capture data. Genome Res. 2017; 27(5):778–86.

24. Sun H, Ding J, Piednoël M, Schneeberger K. FindGSE: estimating genome size variation within human and Arabidopsis using k-mer frequencies. Bioinformatics. 2018; 34(4):550–7.

25. Alonge M, Soyk S, Ramakrishnan S, Wang X, Goodwin S, Sedlazeck FJ, et al. Fast and accurate reference-guided scaffolding of draft genomes. bioRxiv. 2019; :519637.

26. Human Genome Sequencing Consortium I. Finishing the euchromatic sequence of the human genome. Nature. 2004; 431(7011):931–45.

27. The Genome Institute at Washington University School of Medicine. NA12878_prelim_3.0. https://www.ncbi.nlm.nih.gov/assembly/GCA_00207703. 2018.

28. Strope PK, Skelly DA, Kozmin SG, Mahadevan G, Stone EA, Magwene PM, et al. The 100-genomes strains, an S. cerevisiae resource that illuminates its natural phenotypic and genotypic variation and emergence as an opportunistic pathogen. Genome Res. 2015; 25(5):762–74.

29. Goffeau A, Barrell BG, Bussey H, Davis RW, Dujon B, Feldmann H, et al. Life with 6000 Genes. Science (80-). 1996; 274(5287):546–67.

30. Chakraborty M, VanKuren NW, Zhao R, Zhang X, Kalsow S, Emerson JJ. Hidden genetic variation shapes the structure of functional elements in Drosophila. Nat Genet. 2018; 50(1):20–5.

31. Hoskins RA, Carlson JW, Wan KH, Park S, Mendez I, Galle SE, et al. The Release 6 reference sequence of the Drosophila melanogaster genome. Genome Res. 2015; 25(3):445–58.

32. Hirsch CN, Hirsch CD, Brohammer AB, Bowman MJ, Soifer I, Barad O, et al. Draft Assembly of Elite Inbred Line PH207 Provides Insights into Genomic and Transcriptome Diversity in Maize. Plant Cell. 2016; 28(11):2700–14.

33. Jiao Y, Peluso P, Shi J, Liang T, Stitzer MC, Wang B, et al. Improved maize reference genome with single-molecule technologies. Nature. 2017; 546(7659):524.

34. Schnable PS, Ware D, Fulton RS, Stein JC, Wei F, Pasternak S, et al. The B73 maize genome: complexity, diversity, and dynamics. Science. 2009; 326(5956):1112–5.

35. Sun H, Rowan BA, Flood PJ, Brandt R, Fuss J, Hancock AM, et al. Linked-read sequencing of gametes allows efficient genome-wide analysis of meiotic recombination. bioRxiv. 2018; :484022.

36. Zapata L, Ding J, Willing E-M, Hartwig B, Bezdan D, Jiao W-B, et al. Chromosome-level assembly of Arabidopsis thaliana Ler reveals the extent of translocation and inversion polymorphisms. Proc Natl Acad Sci U S A. 2016; 113(28):E4052–60.

37. Rowan BA, Patel V, Weigel D, Schneeberger K. Rapid and Inexpensive Whole-Genome Genotyping-by-Sequencing for Crossover Localization and Fine-Scale Genetic Mapping. G3 Genes, Genomes, Genet. 2015; 5(3):385–98.

38. Imprialou M, Kahles A, Steffen JG, Osborne EJ, Gan X, Lempe J, et al. Genomic rearrangements in Arabidopsis considered as quantitative traits. Genetics. 2017; 205(4):1425–41.

39. Wijnker E, Velikkakam James G, Ding J, Becker F, Klasen JR, Rawat V, et al. The genomic landscape of meiotic crossovers and gene conversions in Arabidopsis thaliana. Elife. 2013; 2.

40. Qi J, Chen Y, Copenhaver GP, Ma H. Detection of genomic variations and DNA polymorphisms and impact on analysis of meiotic recombination and genetic mapping. Proc Natl Acad Sci U S A. 2014; 111(27):10007–12.

41. Paten B, Novak AM, Eizenga JM, Garrison E. Genome graphs and the evolution of genome inference. Genome Res. 2017; 27(5):665–76.

42. Consortium TCP. Computational pan-genomics: status, promises and challenges. Brief Bioinform. 2016; (August):bbw089.

43. Chin C-S, Peluso P, Sedlazeck FJ, Nattestad M, Concepcion GT, Clum A, et al. Phased diploid genome assembly with single-molecule real-time sequencing. Nat Methods. 2016; 13(12):1050–4.

44. Zapata L, Ding J, Willing E-M, Hartwig B, Bezdan D, Jiao W-B, et al. Chromosome-level assembly of Arabidopsis thaliana L er reveals the extent of translocation and inversion polymorphisms. Proc Natl Acad Sci. 2016; 113(28):E4052–60.

45. Smit, AFA, Hubley, R & Green P. RepeatMasker Open-4.0. http://www.repeatmasker.org.

46. Bartenhagen C, Dugas M. RSVSim: an R/Bioconductor package for the simulation of structural variations. Bioinformatics. 2013; 29(13):1679–81.

47. Jeffares DC, Jolly C, Hoti M, Speed D, Shaw L, Rallis C, et al. Transient structural variations have strong effects on quantitative traits and reproductive isolation in fission yeast. Nat Commun. 2017; 8(1):14061.

48. Michael TP, Jupe F, Bemm F, Motley ST, Sandoval JP, Lanz C, et al. High contiguity Arabidopsis thaliana genome assembly with a single nanopore flow cell. Nat Commun. 2018; 9(1):541.

49. Li H. Wgsim: Reads simulator. https://github.com/lh3/wgsim.

50. Li H, Handsaker B, Wysoker A, Fennell T, Ruan J, Homer N, et al. The Sequence Alignment/Map format and SAMtools. Bioinformatics. 2009; 25(16):2078–9.

51. Kielbasa SM, Wan R, Sato K, Horton P, Frith MC. Adaptive seeds tame genomic sequence comparison. Genome Res. 2011; 21(3):487–93.

52. Faust GG, Hall IM. SAMBLASTER: fast duplicate marking and structural variant read extraction. Bioinformatics. 2014; 30(17):2503–5.

53. Ossowski S, Schneeberger K, Clark RM, Lanz C, Warthmann N, Weigel D. Sequencing of natural strains of Arabidopsis thaliana with short reads. Genome Res. 2008; 18(12):2024–33.

54. Benjamini Y, Hochberg Y. Controlling the false discovery rate: a practical and powerful approach to multiple testing. J R Stat Soc Ser B. 1995; 57(1):289–300.

55. Initiative TAG. Analysis of the genome sequence of the flowering plant Arabidopsis thaliana. Nature. 2000; 408(6814):796–815.

56. Goel M, Schneeberger K. SyRI: identification of syntenic and rearranged regions from whole-genome assemblies. https://schneebergerlab.github.io/syri/.

