## Additional File 1 for "SyRI: finding genomic rearrangements and local sequence differences from whole-genome assemblies"

**Supplementary Information**

*Manish Goel, Hequan Sun, Wen-Biao Jiao,* *Korbinian Schneeberger^*^*

Max Planck Institute for Plant Breeding Research, D-50829 Cologne, Germany

^*^ To whom correspondence should be addressed

### Additional Notes

#### Note 1: SyRI input file format

SyRI takes pairwise whole-genome alignments as input. The alignments need to be in a tab-delimited file format where each row corresponds to an alignment and the columns are:

| genome 1 start position (1-based. Includes the start position.) | int |
| --- | --- |
| genome 1 end position (1-based. Includes the end position.) | int |
| genome 2 start position (1-based. Includes the start position.) | int |
| genome 2 end position (1-based. Includes the end position.) | int |
| alignment length on genome 1 | int |
| alignment length on genome 2 | int |
| alignment identity | float |
| alignment direction in genome 1 | int |
| alignment direction in genome 2 | int |
| chromosome ID in genome 1 | string |
| chromosome ID in genome 2. | string |
| CIGAR string (optional) | string |

The rows (alignments) should be sorted based on the alignment start position in genome 1. For genomic difference identification, genome 1 would be considered as the reference genome.

#### Note 2: Synteny identification

For identifying syntenic regions, SyRI selects directed alignments between a pair of homologous chromosomes from the whole-genome alignments between two chromosome-level genome assemblies (Additional File 1: Figure S1). Using these alignments, it generates a directed acyclic graph (DAG). In this DAG, each node corresponds to an alignment, and an edge exists between two nodes when the corresponding aligned regions for these nodes have the same relative position in both genomes. For the addition of edges between two nodes, the corresponding aligned regions do not need to be closely-linked and there should be no other alignment in between these aligned regions on both genomes. Consequently, in Additional File 1: Figure S1, node “a” has an edge to both node “e” and “c” as their corresponding aligned regions are downstream to aligned regions of “a” in both genomes. However, nodes with another alignment between their aligned regions (“c” between “a” and “d”), nodes for which the aligned regions have different order in the two genomes (inter-crossing alignments, “d” and “e”), and nodes from overlapping alignments (“a” and “b”) are not connected by an edge. Each node gets a score equal to the corresponding alignment score. Additionally, two 0-score start (S) and end (E) nodes are also added. Edges are added from node S to all nodes without any in-edge, and from all nodes without any out-edge to node E. Finally, using dynamic programming, the longest path between S 🡪 E is found. Nodes in this path correspond to the syntenic path and the respective alignments are selected as syntenic alignments (alignments with blue boundary).

#### Note 3: Inversion Identification

Alignments from a putative inversion can have different conformations relative to each other resulting in different types of inversion (Additional File 1: Figure S2). Further, these inverted alignments can overlap with each other, which can result in multiple contradicting annotations for an inverted region. From these convoluted inverted alignments, SyRI tries to find alignments which can best annotate these inverted regions while not being restricted by the complexity of the inverted region and thus identifying all types of inversions (Additional File 1: Figure S2).

The inversion identification algorithm is based on the observation that similar to synteny, inversions are also aligning same loci in the two genomes. Correspondingly, if the query genome is reverse complemented, then the alignments corresponding to one inversion will become directed as well as syntenic to each other. These locally syntenic alignments between the reference and reverse complemented query genome correspond to inverted alignments which together can annotate an inversion.

For identifying all inversions between a pair of homologous chromosomes, SyRI first reverse compliments the query chromosome (Additional File 1: Figure S3a). Then it creates a directed acyclic graph (DAG) using the now directed alignments (originally inverted alignments) between the two chromosomes. In the DAG, each node corresponds to an alignment (directed) and an edge is added between two nodes when the relative position of their aligned regions is the same in reference and the reversed query genomes. Similar to synteny identification, for the addition of an edge the corresponding aligned regions of the node do not need to be closely-linked and there should be no other alignment in between these aligned regions on both genomes. Therefore, in Additional File 1: Figure S3a, an edge exists between “a” and “c” because aligned regions for “c” are downstream to aligned regions of “a” on both (reference and reversed query) genomes. However, “a” and “b” do not have an edge because the alignments are inter-crossing. Two 0-score start (S) and end (E) nodes are added and edges are added from node S to all nodes and from all nodes to node E. In this DAG, each S 🡪 E path corresponds to alignments which together can represent an inverted region. The annotation provided by these alignments is termed as candidate inversions.

In Additional File 1: Figure S3b, seven candidate inversions could be identified. These candidates can overlap (ex: candidate 1 and 4) or could be conflicting (ex: candidate 1 and 3 are conflicting because inversions cannot cross each other). To resolve these conflicts, SyRI generates a second DAG where each candidate is a node and an edge exists two nodes when they are non-overlapping, non-conflicting, and no other candidate is between them. Nodes have a score equal to the difference of the sum of its constituent alignments’ score and the sum of alignment score of any syntenic region within the inverted region (since syntenic regions cannot exist inside inverted regions and would need to be removed). Imaginary, start (S) and end (E) nodes are also added. SyRI then adds edges from node S to all nodes without any in-edge, and from all nodes without any out-edge to node E. Using dynamic programming, longest S🡪E path is identified. This path contains all (candidate) inversions which can co-exist while maximizing alignment score for the entire chromosome.

#### Note 4: Details on TD identification

The TD identification starts by selecting all currently un-annotated alignments. These alignments are annotated as either translocations or duplications, or filtered out as redundant (repetitive). Alignments from translocated or duplicated regions can cross syntenic or inverted regions as well as each other (Additional File 1: Figure S4). Additionally, the presence of local sequence differences can result in alignment breaks, resulting in two or more alignments representing one rearrangement (TD) (Additionally File 1: Figure S5a).

SyRI generates a directed acyclic graph (DAG) (Additional File 1: Figure S6) from all un-assigned (i.e. non-syntenic and not included in inversions) alignments. Similar to syntenic path identification, the nodes of the DAG correspond to the alignments (with node score equal to the alignment score) and an edge is added between two nodes when their corresponding aligned regions have same relative position in both genomes and no other single alignment is in between the aligned regions of these nodes in both genomes (e.g. edge between “a” and “b” in Additional File 1: Figure S6). For this, the aligned regions do not need to be closely linked. However, edges are not added between nodes for which the aligned regions are separated by a common syntenic (or inverted) region on both regions (eg. “b” and “d”) or for which the alignments are inter-crossing (eg. “c” and “d”). Imaginary start (S) and end (E) nodes are added and edges are added from the start node to all other nodes and from all other nodes to the end node. In this graph, the alignments of each S🡪E path represent a TD rearrangement.

However, similar to the overlapping of individual alignments, these alignment groups can overlap too resulting in conflicting annotations (Additional File 1: Figure S5b). Therefore, each of these identified group is only a candidate for explaining a putative genomic rearrangement and is therefore termed as candidate TD. SyRI scores each candidate based on the alignments lengths and the gaps between its consecutive alignments; score $\boldsymbol{=}min \left( \frac{genA\_aligned\_length-genA\_gap\_length}{genA\_aligned\_length}, \frac{genB\_aligned\_length-genB\_gap\_length}{genB\_aligned\_length} \right)$. All candidates with a negative score (i.e. having large gaps between the alignments) are filtered out.

Since rearranged regions are often rich with repeats, they can have multiple copies in the two genomes. The different copies from a genome can align to different copies in the other genome, leading to multiple overlapping candidates forming a network of candidate TDs (Additional File 1: Figure S5c). Such a network consists of all candidates which overlap each other or any other candidates in the network. For example, in Additional File 1: Figure S5c, a blue candidate is overlapping with two green candidates thus forming a network. Since the other blue candidate is also overlapping with the green candidates, it will be also added to the same network. Therefore, even though the two blue candidates are not overlapping with each other, they are still part of the same candidate network of. SyRI generates this network by first selecting a random candidate as the *seed* candidate for the network. Then, all candidates overlapping (at either of the two genomes) this seed candidate are also selected as members of this network. Then, candidates overlapping these new member candidates are also added to this network. This process is repeated until no other overlapping candidate can be added.

In a network, the annotation for a candidate TD would depend on the annotation of its overlapping candidates. For example, in Additional File 1: Figure S5c, for the first case the two larger green candidates would be selected as two translocations while the smaller cyan candidates would be removed as redundant. However, in the second case, the cyan candidates are larger than green candidates and explain the genomic differences better, hence these would be selected as translocations and the green candidates would be removed as redundant. Thus, the annotation for a candidate can change even without any changes in the candidate itself. Consequently, for accurate annotation of genomic rearrangements, it is important to consider the inter-dependence of all candidates in a network by performing a comparative selection of these candidates simultaneously.

SyRI compares all candidates in a network and selects candidates which together provide the best annotation for the genomic rearrangements. As the first step, SyRI selects candidates which align to a region uniquely (i.e. candidates for which the aligned region does not overlap with other candidates or syntenic/inverted region) (Additional File 1: Figure S7a). These *unique candidates* are necessary for annotating corresponding genomic regions as there are no alternative candidates that could annotate these regions. Therefore, these are selected as part of the final group of selected candidate TDs. Then, candidates aligning already annotated regions within both genomes are filtered out as redundant (repetitive) candidates. The process of selection of unique (necessary) candidates and removal of redundant (unnecessary) candidates is termed as *progressive elimination* and is performed iteratively.

However, it is possible that none of the remaining candidates in a network could either be selected as unique or removed as redundant (Additional File 1: Figure S7b). This is termed as a *deadlock* as it is not possible to continue with progressive elimination anymore. SyRI resolves such deadlocks by using a brute-force approach for smaller networks and a randomized-greedy approach for larger networks.

In the brute-force approach, SyRI enumerates all combinations in which candidates in a network can be grouped (Additional File 1: Figure S7c). From this list of groups, it selects the highest-scoring group and then its member candidates are used for annotating genomic rearrangements. For this, SyRI starts by selecting all uniquely aligning candidates as the *seed* group, thus initiating the list of groups. Then, it iterates over the list of all remaining candidates in the network, and if a candidate can co-exist with the candidates of a group in the group-list, then a new group consisting of the current candidate and the candidates in the group is added to the list of groups. If the number of different groups becomes too large, SyRI automatically switches to a randomized-greedy algorithm to ensure limited memory usage and lower run-time.

In the randomized-greedy approach, SyRI randomly selects one of the highest-scoring candidates to break the deadlock after which it continues with progressive elimination. Like earlier, all uniquely aligning candidates are selected as the *seed* group. Then to resolve the deadlock, SyRI selects one of the twenty highest-scoring candidates and adds it to the group. The selection probability of a candidate is proportional to its score. Addition of a candidate breaks the deadlock after which SyRI again starts removal of redundant candidates and selection of uniquely aligning candidates. This process is continued until all candidates in the network are either selected or filtered out, resulting in a group of selected candidates which can co-exist. Starting from the seed group, this process is repeated 100 times, resulting in 100 groups of candidate TDs.

For both brute-force and randomized-greedy approach, each group is given a score equal to the number of new (previously unannotated) bases its constituent candidates annotates. SyRI uses this score to select candidate groups which best explain genomic rearrangements. Additionally, while selecting the best group, SyRI gives preference to groups comprising of few longer candidates compared to groups with a similar score but consisting of multiple small candidates.

SyRI then classifies the candidates of the best group as either translocations or duplications based on their location in the genomes. For this, it checks the overlap between selected candidates and the already annotated syntenic/inverted regions. If a candidate does not overlap with already annotated regions or with other selected candidates, then it is classified as translocation. Whereas, if the candidate overlaps with a syntenic/inverted region then the candidate is considered as a duplication. When two candidates overlap each other on one genome but on the other genome align uniquely (i.e. do not overlap each other or any other annotated region), then the candidate with the higher score is selected as translocation while the other is annotated as duplication.

#### Note 5: Pseudo-genome generation

To apply SyRI to assemblies, which are not at chromosome-level, pseudo-chromosomes can be generated using homology to a chromosome-level assembly. Typically, this will lead to false negative predictions if certain rearrangements were not present in the original assembly, however, usually, this will not lead to an increased rate of false positives. To generate such assemblies, one can use dedicated tools like RaGOO (preprint [1]) and Chromosomer [2]. These tools generate pseudo-chromosomes by concatenating scaffolds of one assembly based on a chromosome-level assembly.

However, if both assemblies are at scaffold-level, most of these tools cannot be applied. For such cases, we developed a heuristic method which uses whole genome alignments between the two scattered assemblies to generate ordered sets of contigs that can be matched between the two assemblies and that can be used as pseudo-chromosomes when running SyRI. The method finds homologous scaffolds and then orders and orients them based on their alignments with the corresponding homologous scaffold. When a scaffold in one assembly aligns with multiple scaffolds in the other assembly then these scaffolds are selected as constituents to form a larger pseudo-chromosome. The order in which these scaffolds need to be arranged is determined based on their respective alignment location in other scaffolds, whereas alignment direction is used to get scaffold orientation. A scaffold is assigned to only one chromosome and no breaks are introduced. Scaffolds which could not be uniquely assigned to a chromosome are filtered out. Finally, scaffolds which form linear sequence are ordered and concatenated with Ns between adjacent scaffolds.

#### Note 6: Optimal rearrangement identification is highly complex

SyRI tries to identify the optimal set of structural rearrangements. However, non-syntenic alignments, which will constitute structural rearrangements, overlap each other. Consequently, these alignments and the corresponding rearrangements can result in conflicting annotations such that only one of many rearrangements can be selected. For such alignments, SyRI tries to list all different rearrangements that can be generated using these alignments and then selects rearrangements which would lead to highest scoring annotation (Additional File 1: Figure S19).

### References

1.	Alonge M, Soyk S, Ramakrishnan S, Wang X, Goodwin S, Sedlazeck FJ, et al. Fast and accurate reference-guided scaffolding of draft genomes. bioRxiv. 2019; :519637.

2.	Tamazian G, Dobrynin P, Krasheninnikova K, Komissarov A, Koepfli K-P, O’Brien SJ. Chromosomer: a reference-based genome arrangement tool for producing draft chromosome sequences. Gigascience. 2016; 5(1):38.

### Supplemental Figures

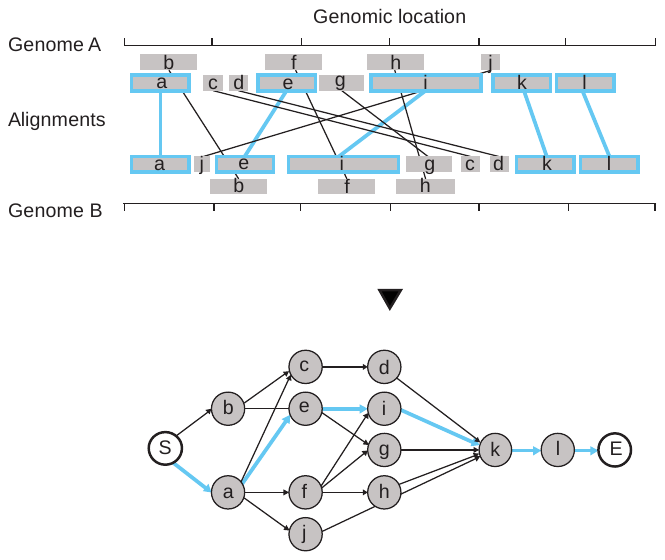

Figure S1: Syntenic path identification. Alignments between two genomes (Genome A and Genome B) are shown by grey blocks (top). Overlapping alignments are stacked vertically (“b” is overlapping “a” on Genome A and it is overlapping “e” on Genome B). A directed acyclic graph (DAG) is formed using these alignments (bottom). The grey nodes correspond to the alignments, while white nodes (“S” and “E”) are 0-score imaginary nodes. The blue line in the graph represents the identified syntenic path and the blue boundary on alignments are the corresponding alignments.

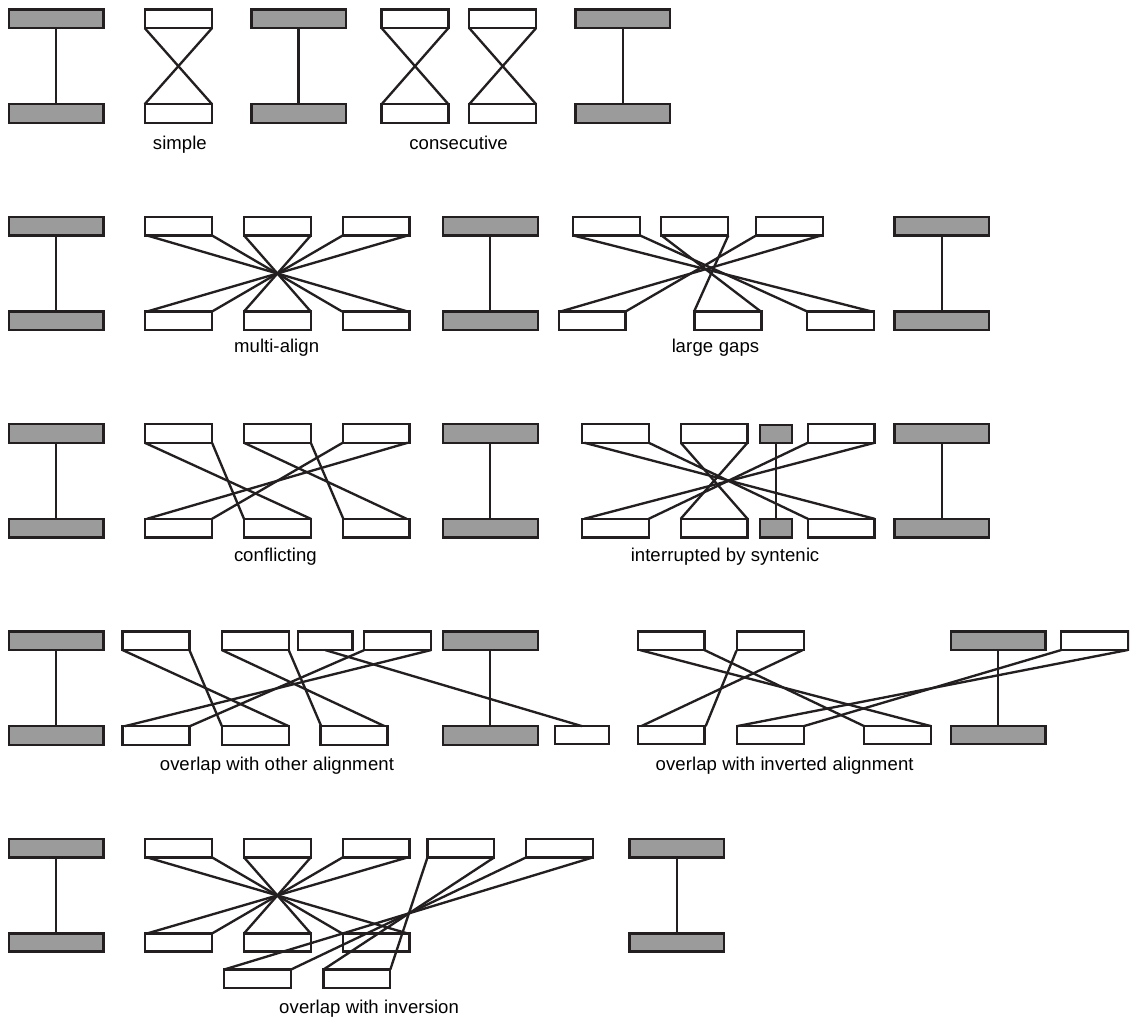

Figure S2: Different types of inversions which can be identified by SyRI. Inversions comprise of inverted alignments (white alignments with dashed lines) surrounded by syntenic alignments (grey alignment) on both sides. In the easiest case, inversions can consist of a *simple* inverted alignment. Alternatively, *multiple inverted alignments* can correspond to a single inversion event, or multiple inversion events could occur *consecutively*. Also, a region can show signals for multiple *conflicting* inversions. Alignments corresponding to inversions can be *overlapping* or *interrupted* by TDs or syntenic regions.

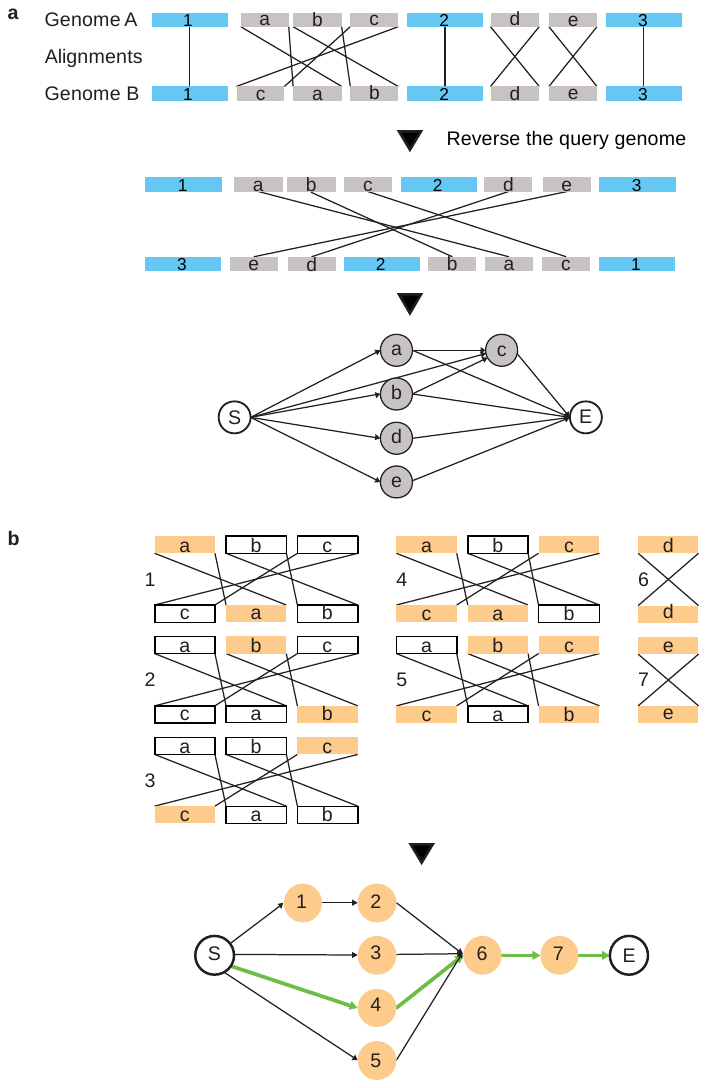

Figure S3: Inversion identification. (a) Inverted alignments between two genomes (Genome A and Genome B) are shown by grey blocks (top), blue blocks represent the already annotated syntenic regions. A directed acyclic graph (DAG) is formed using these alignments (bottom). The grey nodes correspond to the alignments, while white nodes (“S” and “E”) are 0-score imaginary nodes. Each S🡪E path in this graph correspond to a candidate inversion. (b) Seven different candidate inversions are found. Orange blocks represent inverted alignments which constitute the candidate inversion. A directed acyclic graph (DAG) is formed using these candidates. The orange nodes correspond to the candidates, while white nodes (“S” and “E”) are 0-score imaginary nodes. The green line in the graph represents the highest scoring path from node S to node E.

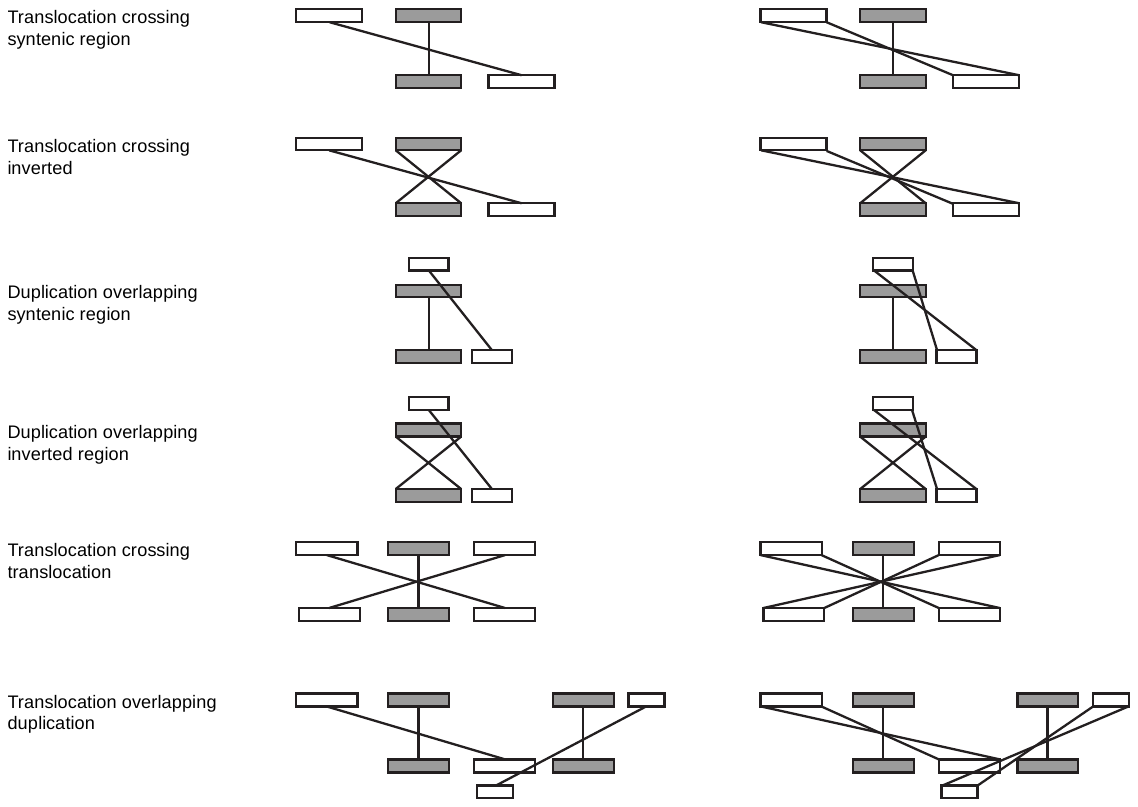

Figure S4: Different types of translocations and duplications (TDs). TD alignments (white) cross or overlap already annotated (grey) alignments and each other. TD alignments can be directed or inverted.

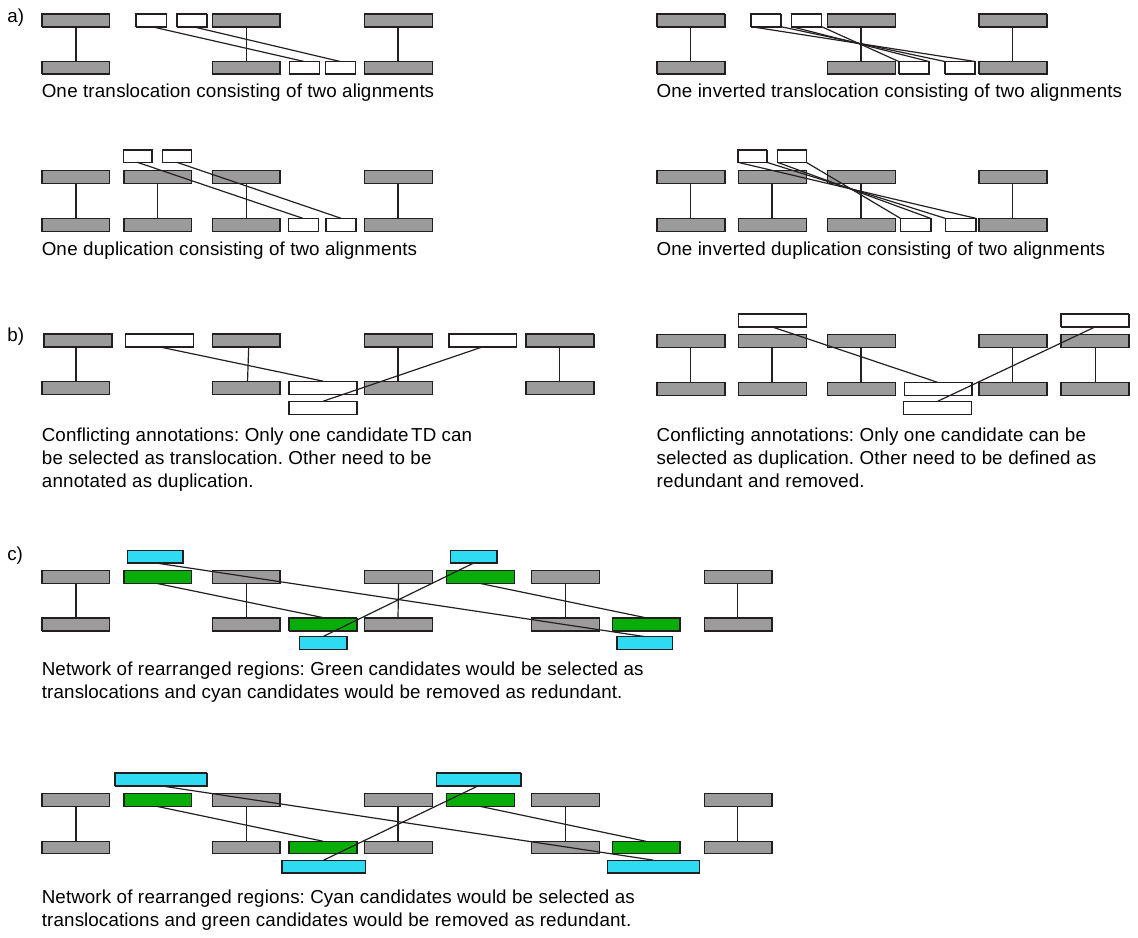

Figure S5: Complexities of translocations and duplication identification. The grey blocks represent syntenic/inverted regions. (a) White blocks are alignments (directed or inverted) between rearranged regions. Local sequence differences in rearranged regions can result in alignment breaks. Consequently, one TD event could consist of multiple alignments. These different alignments are grouped and selected as candidate TD. (b) White blocks represent candidate TDs. The candidate TDs can overlap each other resulting in conflicting annotations. In this case, the rearranged region in the lower genome can be annotated as translocation (or duplication) from two different regions. Since one region cannot be translocated (or duplicated) from two regions, it is necessary to select candidates which best explain the rearrangements. (c) Candidate TDs (in green and cyan blocks) form a network of overlapping candidates. The annotation of a candidate depends on other candidates in the network. In the first case, the green candidates are larger and represent the genome differences better so they would be selected as translocations and the cyan candidates would be removed as redundant. In the second case, the same green candidates would be removed as redundant as the cyan candidates now are larger and represent the genomic differences better.

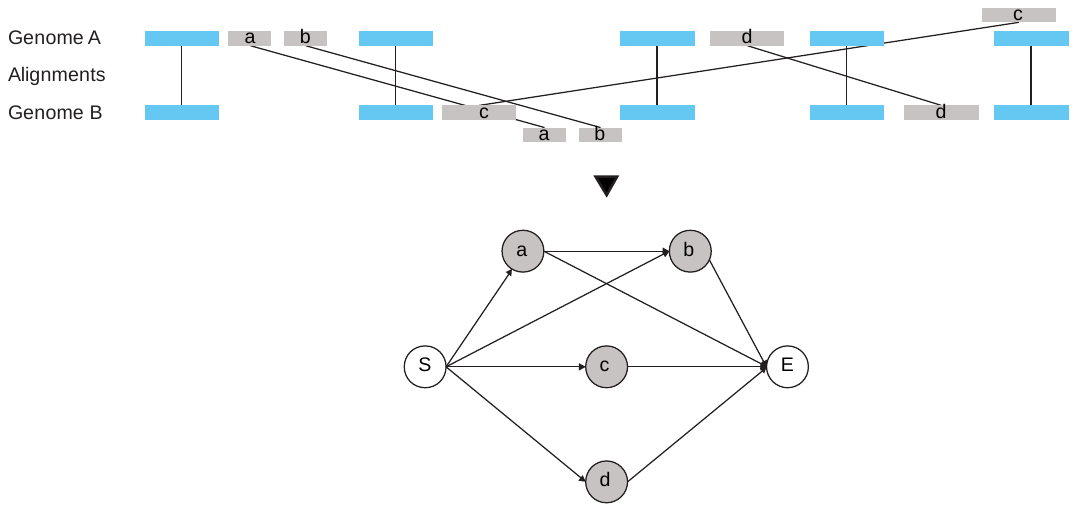

Figure S6: Candidate TD identification. All currently un-annotated alignments (grey blocks) are selected. Blue blocks are alignments between annotated syntenic/inverted regions. A directed acyclic graph is formed where each node (grey) represents an alignment and edges are added between nodes which can be part of a TD rearrangement. Two 0-score nodes, start (S) and end (E), are also added (white nodes) and edges are added from node S to all other nodes and from all other nodes to node E. Each S🡪E path corresponds to a candidate TD.

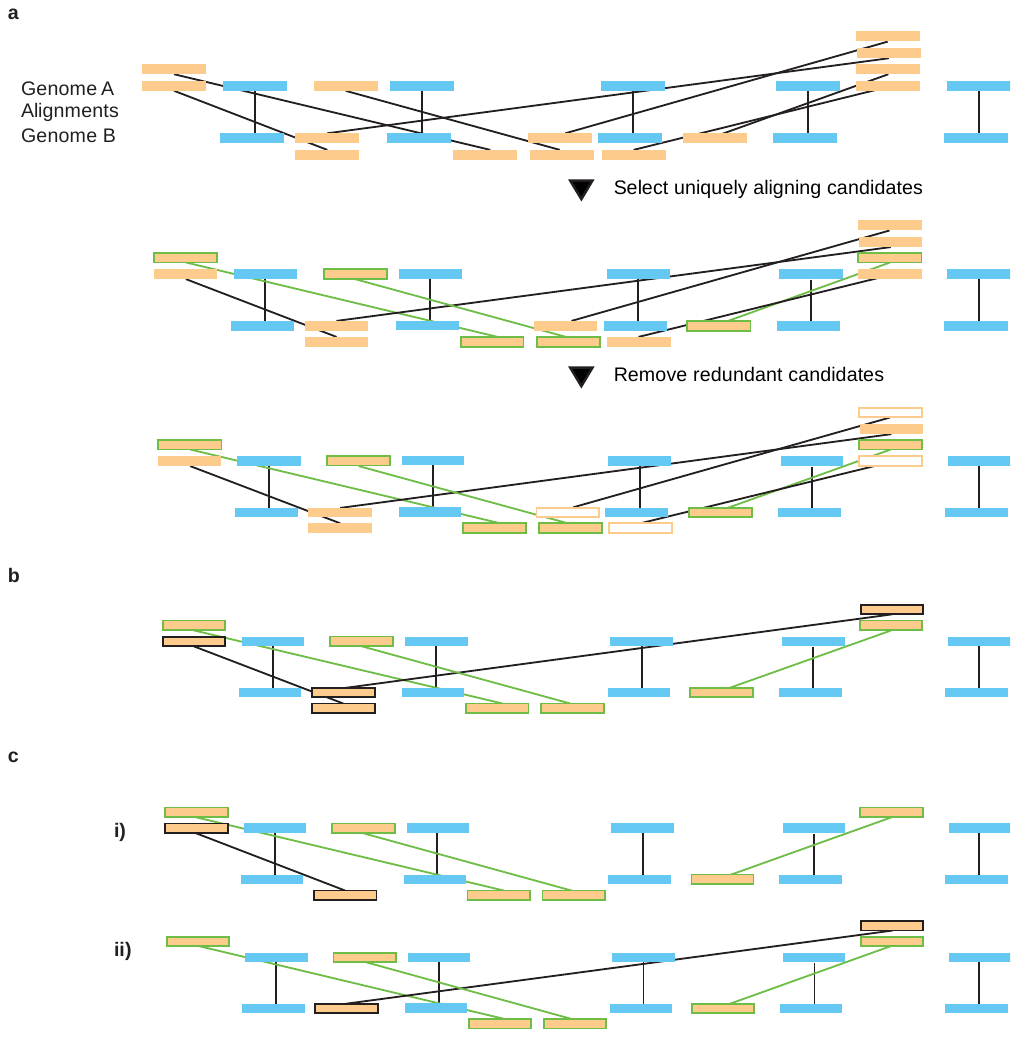

Figure S7: Resolving conflicts and selecting candidate TDs. Candidate TDs (orange alignments) can overlap each other and syntenic/inverted regions (blue alignments) forming a network of co-aligned regions. In a), one such network is shown where three regions on genome A have candidate TD alignments to five regions on genome B. SyRI selects all candidates which align uniquely (green boundary candidates) to a region in either of the two genomes. These uniquely aligning candidates are necessary to annotate the uniquely aligned regions and are therefore selected as part of the output group of candidates. Next, candidates which align already annotated regions (white candidates) are removed as redundant candidates. b) Occasionally, none of the remaining candidates can either be selected as necessary (uniquely aligning) or removed as redundant (black boundary candidates). This is called a deadlock. For networks with fewer candidates SyRI resolves deadlocks by using brute-force approach (c) in which it lists all possible groups in which the remaining candidates can be selected (group i) and ii)) and selects the group which gives the highest alignment score. For networks with a large number of candidates, SyRI randomly selects one of the high scoring candidates as part of the output group. This breaks the deadlock and then the method continues as described in (a).

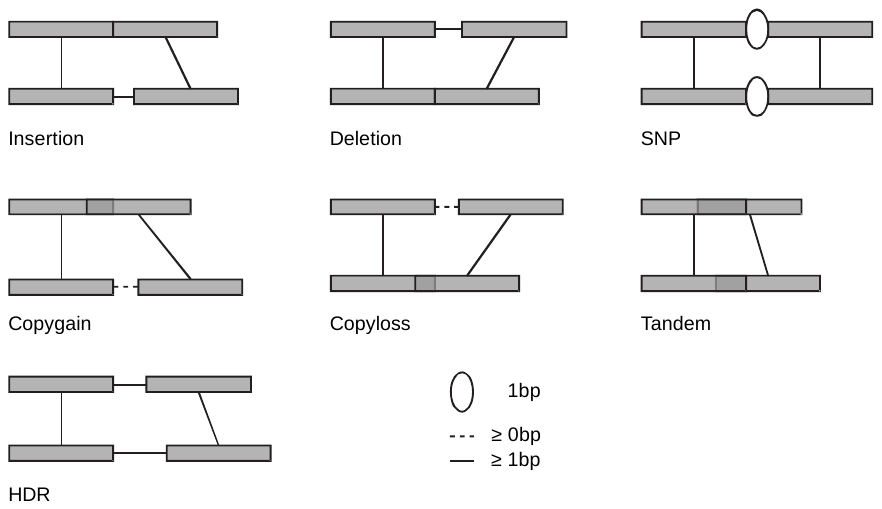

Figure S8: Structural variations identified by SyRI. The arrangement of end-points of consecutive alignments in an annotation block (grey alignments) is analysed to find structural variations.

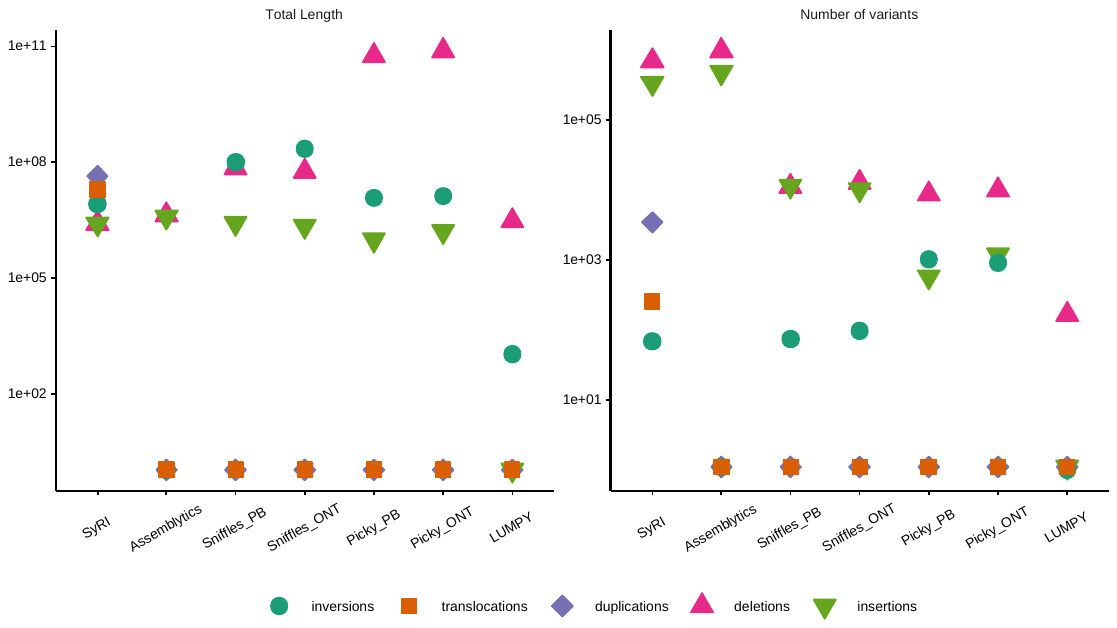

Figure S9: Total length (in bp) and number of structural rearrangements and indels identified by different tools in the NA19240 genome with respect to the human reference genome. Smartie-sv gave no output when run using real genome assemblies and hence could not be analysed. Whereas running AsmVar was too computationally challenging for the human genome and therefore it could not be analysed.

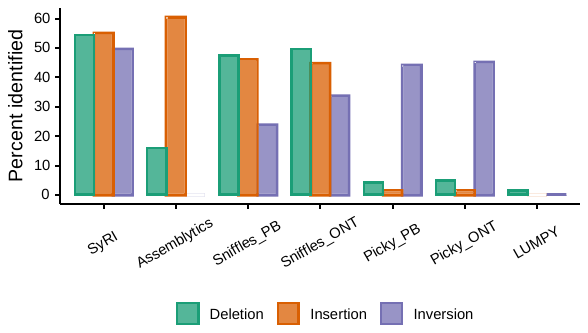

Figure S10: Ratio of gold-standard genomic differences identified by different variant callers. Smartie-sv gave no output when run using real genome assemblies and hence could not be analysed. Whereas running AsmVar was too computationally challenging for the human genome and therefore it could not be analysed.

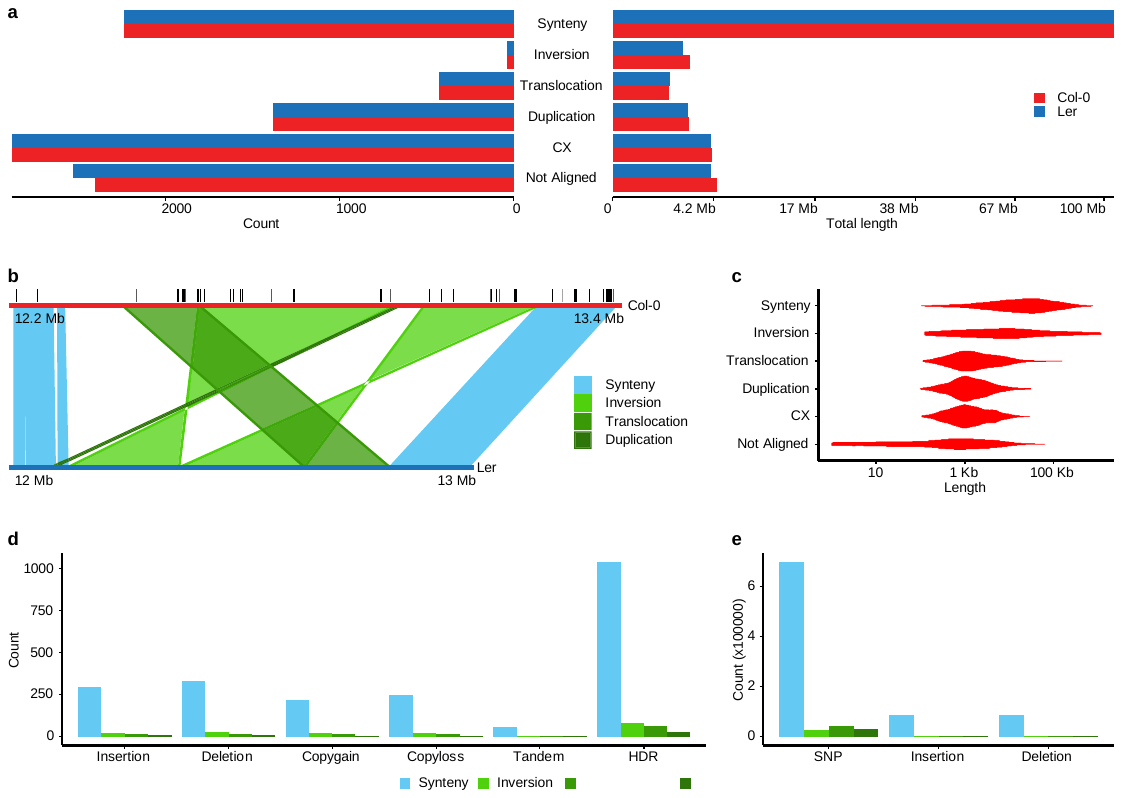

Figure S11 Genomic differences between A. thaliana Col-0 and L*er.* (a) Total length and number of syntenic, structurally rearranged, and not-aligned regions identified between the Col-0 and L*er* genomes (CX: cross-chromosomal exchange). (b) Example of a highly rearranged region at chromosome 5. Two large inversions, a translocation, and a duplication occur together. Multiple genes (black lines) are present in this region. Smaller SRs in this region have been filtered for illustrative clarity. (c) Length distribution of all annotation types. (d) The number of structural variations found by comparing the overlaps and gaps between consecutive alignments in annotation blocks. (e) Number of SNPs and small insertions and deletions identified from within alignments.

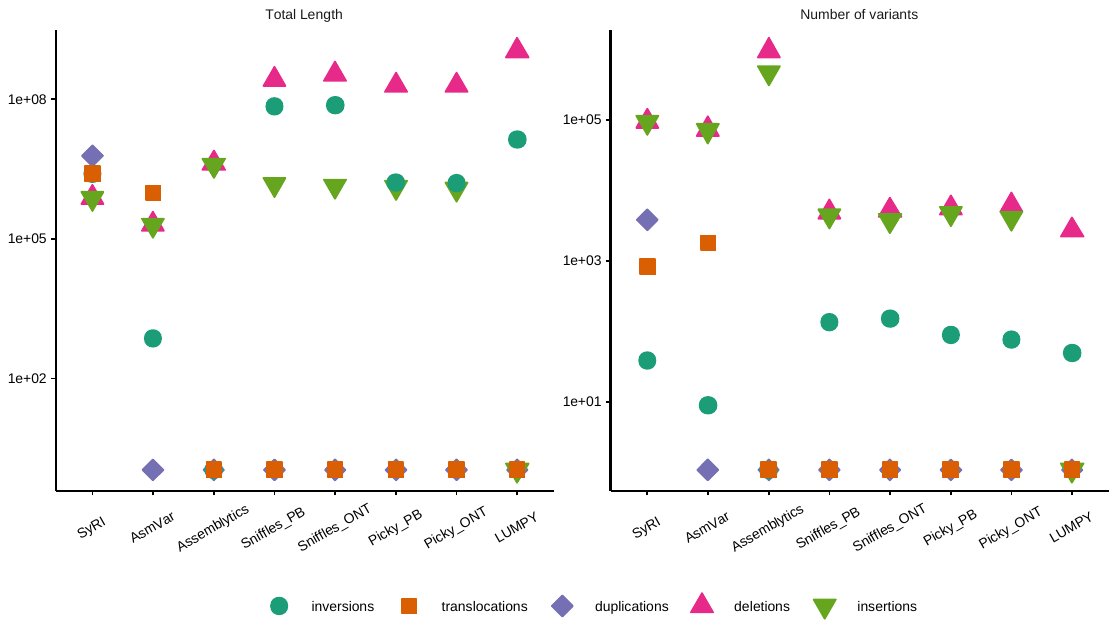

Figure S12: Total length (in bp) and number of structural rearrangements and indels identified by different tools in the L*er* genome with respect to the Col-0 genome. Smartie-sv gave no output when run using real genome assemblies and hence could not be analysed.

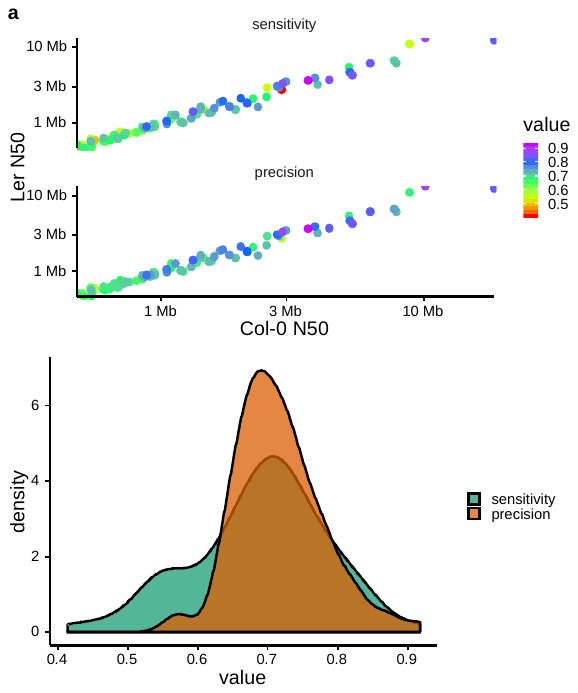

Figure S13: SyRI’s performance statistics when both assemblies are incomplete. (a) Each sample is represented by a point while the colour represents sensitivity and precision values. The x-axis represents N50 values in fragmented Col-0 assembly and the y-axis represents N50 value in fragmented L*er* assembly. (b) Distribution of sensitivity and precision values across analysed 84 samples.

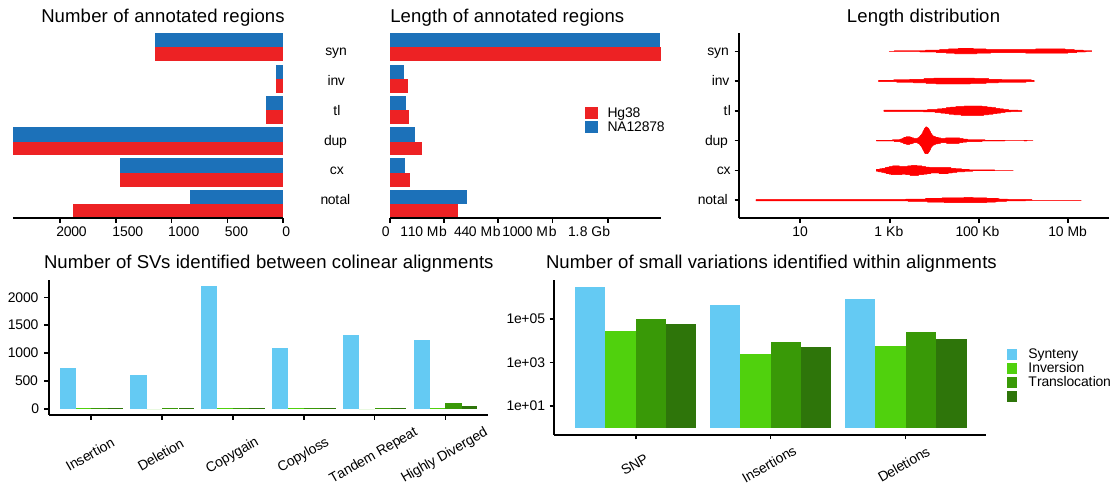

#### Figure S14: Genomic differences identified between human reference and NA12878.

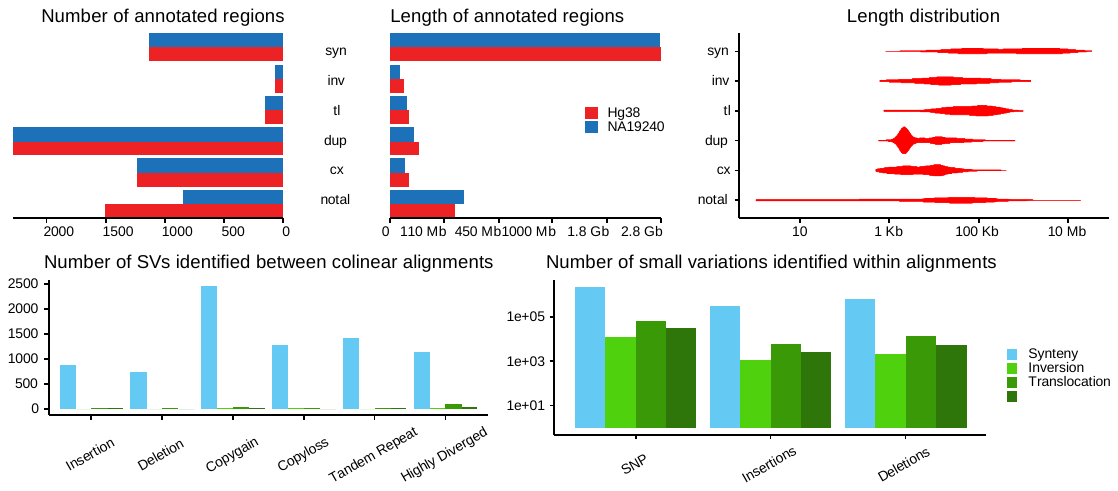

#### Figure S15: Genomic differences identified between human reference and NA19240.

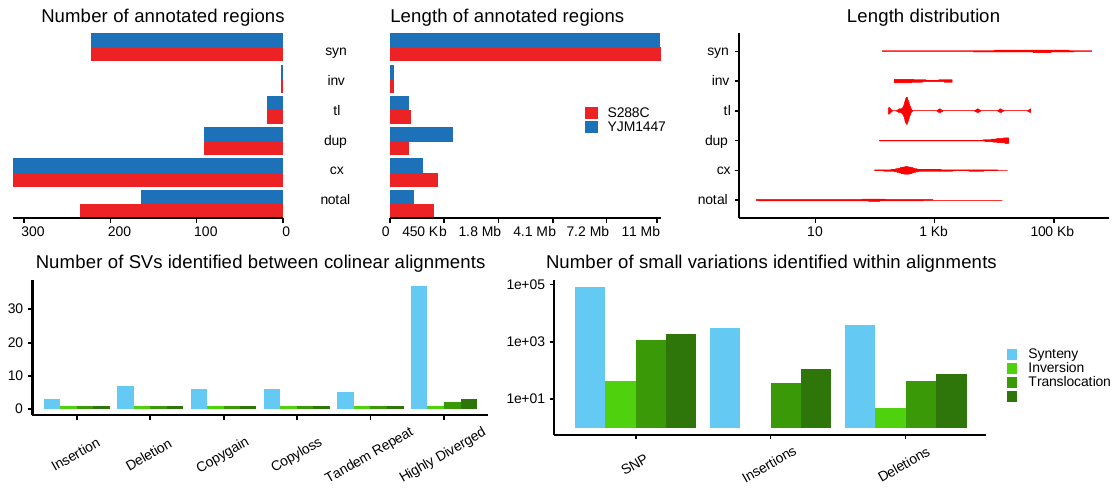

#### Figure S16: Genomic differences identified between yeast reference genome (S288C) and strain YJM1447.

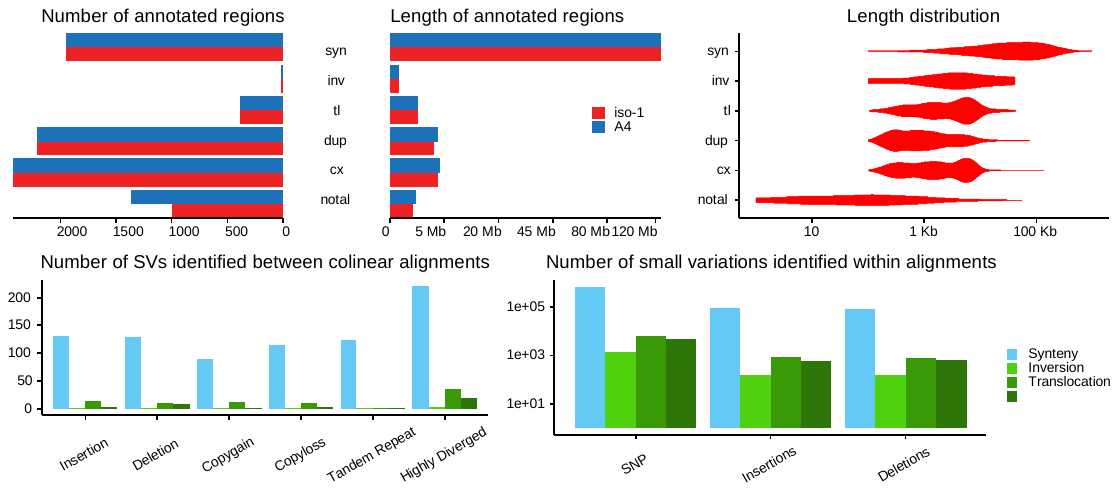

#### Figure S17: Genomic differences identified between fruit fly reference genome (iso-1) and strain A4.

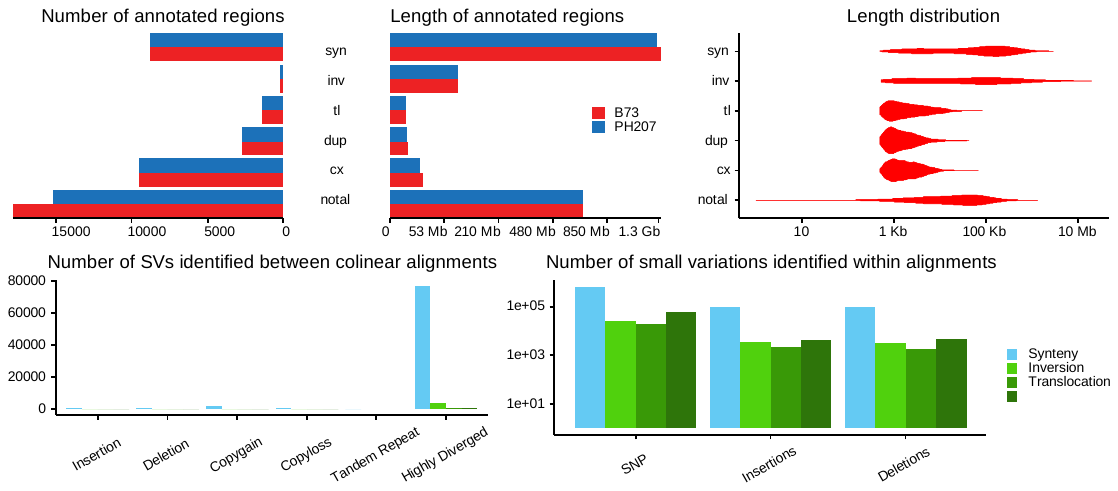

#### Figure S18: Genomic differences identified between the maize reference genome (B73) and strain PH207.

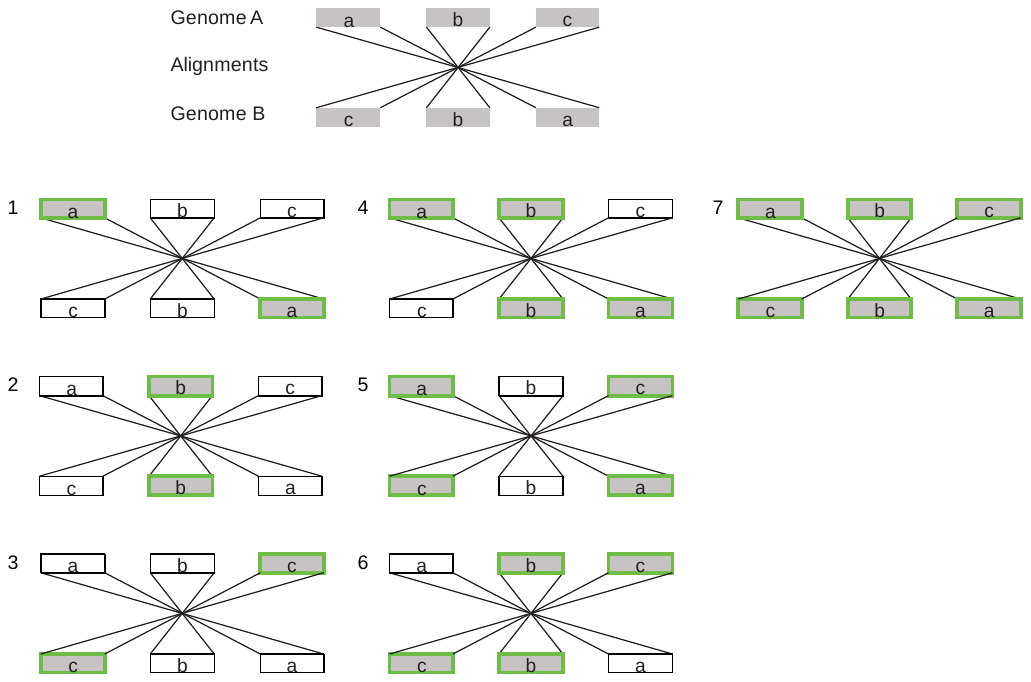

Figure S19: Example of relationships between the number of related alignments and the number of candidate rearrangements. Here, three inverted alignments can correspond to seven candidate inversions (green borders).
